# Enhanced hippocampal theta rhythmicity and emergence of eta oscillation in virtual reality

**DOI:** 10.1101/2020.06.29.178186

**Authors:** Karen Safaryan, Mayank R. Mehta

## Abstract

Hippocampal theta oscillations in rodents profoundly impact neural activity, spatial coding, and synaptic plasticity and learning. What are the sensory mechanisms governing slow oscillations? We hypothesized that the nature of multisensory inputs is a crucial factor in hippocampal rhythmogenesis. We compared the rat hippocampal slow oscillations in the multisensory-rich real world (RW) and in a body-fixed, visual virtual reality (VR). The amplitude and rhythmicity of the hippocampal ~8 Hz theta were enhanced in VR compared to RW. This was accompanied by the emergence of a ~4 Hz oscillation, termed the eta rhythm, evident in the local field potential (LFP) in VR, but not in RW. Similar to theta, eta band amplitude increased with running speed in VR, but not in RW. However, contrary to theta, eta amplitude was highest in the CA1 cell layer, implicating intra-CA1 mechanisms. Consistently, putative CA1 interneurons, but not pyramidal neurons, showed substantially more eta modulation in VR than in RW. These results elucidate the multisensory mechanisms of hippocampal rhythms and the surprising effects of VR on enhancing these rhythms, which has not been achieved pharmacologically and has significant broader implications for VR use in humans.

**One Sentence Summary:** Navigation in virtual reality greatly enhances hippocampal 8Hz theta rhythmicity, and generates a novel, ~4Hz eta rhythm that is localized in the CA1 cell layer and influences interneurons more than pyramidal neurons.

## Main Text

Hippocampal theta oscillations in rodents profoundly impact neural activity (*1, 2*), spatial coding (*3–5*), and synaptic plasticity and learning (*6, 7*). What are the sensory mechanisms governing theta (*8, 9*), and their possible broader implications (*10–12*)? We hypothesized that the nature of multisensory inputs is a crucial factor in hippocampal rhythmogenesis. Hence, we compared the rodent’s hippocampal slow oscillations in the multisensory-rich real world (RW) and in a body-fixed, visual virtual reality (VR) (*13, 14*). Rats were trained to run on a 2.2 m long linear track, either in RW or visually identical VR, to receive rewards at both ends of the track (*14*). LFP was measured from 991 and 1637 dorsal CA1 tetrodes of 4 and 7 rats across 60 RW and 121 VR sessions, respectively.

Consistent with previous studies (*14, 15*), strong 6-10 Hz theta (θ) oscillations were evident in the LFP when the rats were running in either RW or VR (Fig. 1, A and B, top traces, figs. S1 and S2), and were significantly reduced in amplitude at lower speeds (Fig. 1, A and B, bottom traces). However, during runs at higher speeds in VR, but not in RW, clear 2-5 Hz oscillations were also detected on several tetrodes (Fig. 1, A and B, figs. S1 and S2); these are termed hippocampal eta (ƞ) oscillations. Like theta, eta oscillations were significantly enhanced at high compared to low speeds (Fig. 1B). As a result, for many tetrodes, the power spectra during runs in VR revealed a peak not only in the theta (~7.5 Hz), but also in the eta (~4 Hz) band (Fig. 1B) which was absent during immobility. This is in contrast to the power spectra in RW that exhibited a single peak at ~8 Hz during run, as is commonly seen (*14–16*) (Fig. 1A).

**Fig. 1.**
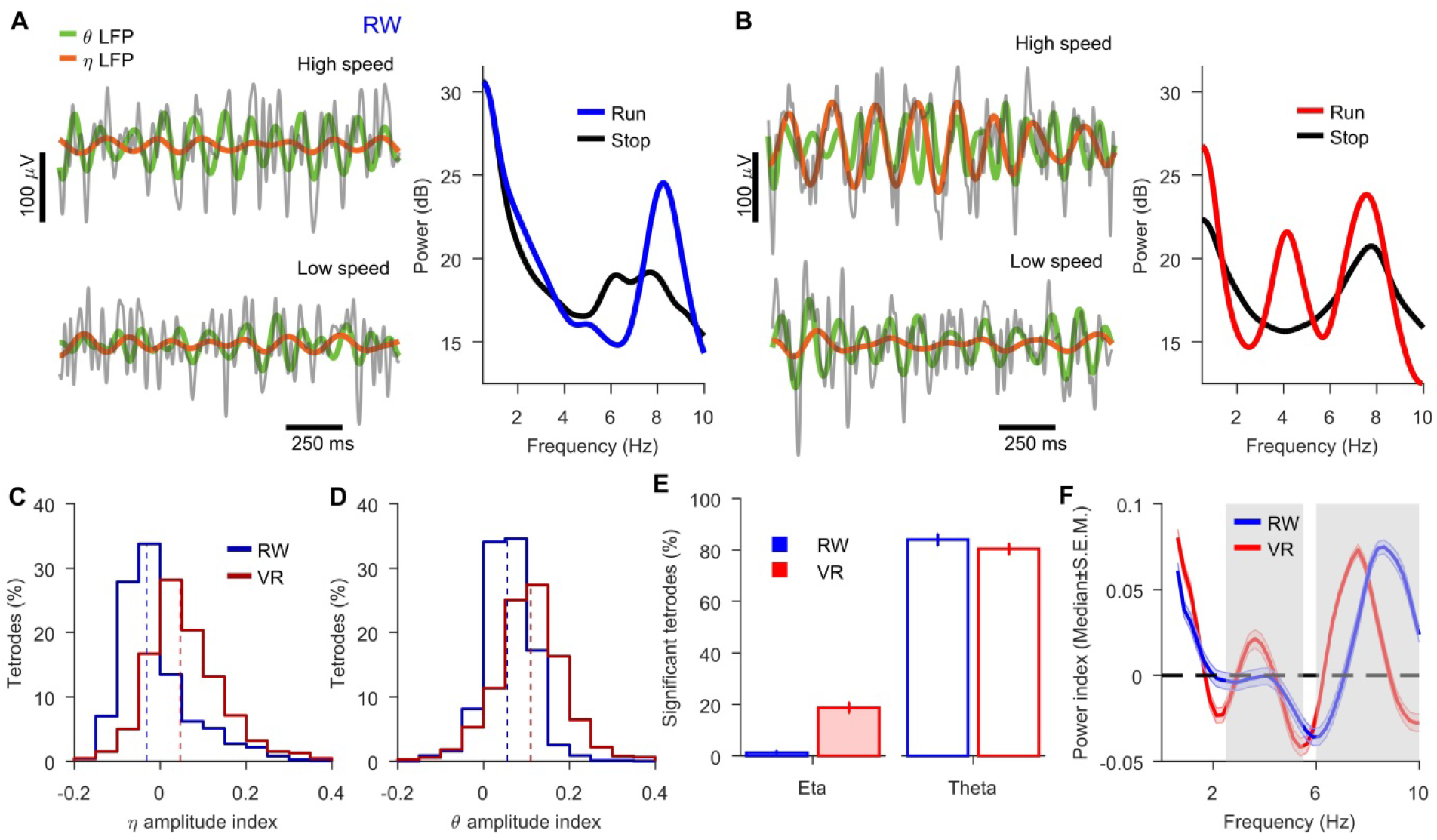
Emergence of a ~4 Hz eta oscillation during running in VR, but not in RW. (A, B) Left, the LFP, raw (grey), filtered in theta (6-10 Hz, green) and eta (2.5-5.5 Hz, brown) bands during high-speed running (> 15 cm/s) on track (top) and at low-speeds (< 15 cm/s, bottom) recorded on the same tetrodes on the same day in the RW (A) and VR (B). Right, power spectra of these LFPs computed during the entire RW (blue) and VR (red) sessions at high- and low-(including stops) speeds. (C) Distributions of the eta amplitude index (see methods) across electrodes in VR (0.063±0.002, red) was significantly greater (p<10^−10^, χ^2^ = 540.5) than in RW (−0.013 ± 0.003, blue). (D) Similar to C, but for theta. Theta amplitude index in VR (0.115 ± 0.001, red) was significantly greater (p<10^−10^, χ^2^ = 414.9) than in RW (0.056 ± 0.002, blue). (E) Left: 1.1% (n = 11, 95% confidence interval [0.6% 1.9%]) and 18.6% (n = 304, binomial test, 95% confidence interval [16.8% 20.6%]) of all recorded tetrodes showed significant, sustained eta oscillatory activity during runs in RW and VR, respectively. Right: 84.1% (n = 821, 95% confidence interval [81.6% 86.2%]) and 80.4% (n = 1310, 95% confidence interval [78.4% 82.3%]) of all tetrodes showed significant, sustained theta oscillatory activity during runs in RW and VR, respectively. (F) Population averaged power index (between run and stop, see methods) for tetrodes recorded on the same day in RW and VR (n = 150, obtained from 4 rats and 39 sessions) that showed significant, sustained eta in RW or VR.

How can these observations be quantified across a variety of RW and VR sessions? The LFP spectral power could be influenced by several nonspecific factors, e.g. the electrode impedance, anatomical localization, and behavioral variability. Furthermore, hippocampal type 2 theta (4-6 Hz) (*17–19*) occurs during periods of immobility in RW. Hence, we computed the LFP amplitude difference between periods of high (30-60 cm/s) and low (5-15 cm/s) speed runs in RW and VR. Remarkably, 71% of all tetrodes in VR showed significantly greater eta amplitude during high speed running epochs, compared to only 29.9% in RW. The eta amplitude contrast index (LFP eta amplitude difference between high and low speed runs divided by their sum or amplitude index) was (600%) greater in VR than in RW (Fig. 1C). The theta amplitude index was also (100%) greater in VR than in RW (Fig. 1D) which is different from previous reports (*14, 15*), where differences in running speeds were not weighted.

One reason for the enhanced theta could be that we quantified the difference in theta (and eta) band amplitudes at high versus low speeds. It is possible that these are occurring in short bouts, only intermittently and not in a sustained fashion. Hence, we examined LFP power spectra in windows of 4s. long epochs, during running and immobility states separately. Similar to the amplitude index, we computed power index as power difference during run and stop at each frequency, and detected tetrodes with significant, prominent peaks in the eta or theta bands (see methods). Overall, 18.6% of all tetrodes in VR showed significantly prominent eta power index peaks along with an increase of the eta amplitude during high speed runs compared to only 1.1% tetrodes in RW (Fig. 1E). Similar analysis of the theta band revealed comparable power in RW and VR (Fig. 1E). To confirm this further, we restricted the analysis to the LFP data from tetrodes recorded in both RW and VR experiments on the same day without any intervening tetrode adjustments. Two distinct peaks in eta and theta bands were observed in VR in comparison to only a single peak in theta band in RW (Fig. 1F). These results showed significant and sustained increase in eta oscillations during run, compared to stop, in VR but not in RW. In general, rats tended to run a bit slower and had greater periods of immobility in VR than in RW (fig. S3), which could explain the reduction in estimation of the theta power spectra compared to the theta amplitude. The results provided by the theta amplitude index are more robust to behavioral differences.

Why was significant eta, often seen in simultaneously recorded electrodes, detected only on a subset of the electrodes (fig. S4)? We hypothesized that the anatomical depth of the electrodes in CA1 is a key determining factor. The lowest theta and sharp wave (SPW) amplitudes occur near the CA1 pyramidal cell layer (20-22). Both theta and SPW amplitudes increase away from the cell layer, while SPW polarity reverses at the cell layer. Thus, SPW amplitude and polarity provide an accurate estimate of the anatomical depth of an electrode. Hence, we measured the amplitude and polarity of SPWs during the baseline sessions preceding the tasks, and compared them to the theta or eta power on the same electrodes during run in VR (Fig. 2, see methods). Consistent with previous findings (*20, 22, 23*), SPW amplitude was significantly correlated with the theta power for both the positive and negative polarity SPWs, such that the smallest theta occurred on tetrodes with the smallest SPW (Fig. 2D). In contrast, the eta power during run was significantly anti-correlated with the SPW amplitude for both the positive and negative polarity SPWs, with the highest eta power coinciding with the lowest SPW amplitude (Fig. 2E).

**Fig. 2.**
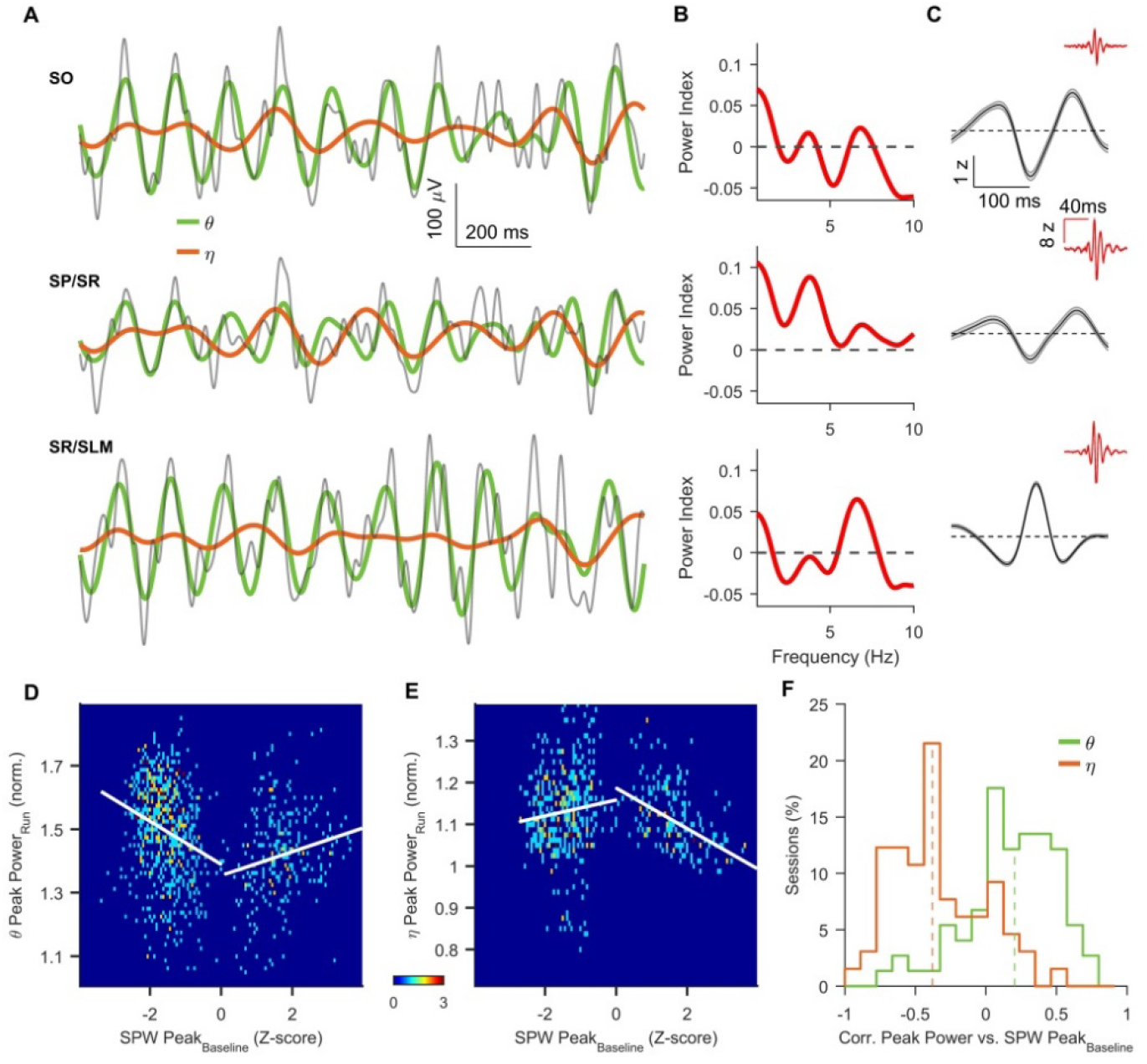
Theta is weakest and eta is strongest in the CA1 cell layer. (A) LFP from three simultaneously recorded tetrodes (same color scheme as Fig. 1A) in a VR session during high-speed (>30 cm/s) run. (B) LFP power index (same as in Fig. 1F) for these electrodes (red). (C) Average z-scored (mean±s.e.m.) ripple traces (red, centered at the peak of the ripple powers) and associated sharp waves (black) for the corresponding electrodes computed during the baseline session preceding the task. The eta band signal (brown) is the highest in the middle row which has the smallest SPW amplitude, while the theta band signal (green) shows the opposite pattern. SO(stratum oriens), SP(stratum pyramidale), SR(stratum radiatum), SLM(stratum lacunosum moleculare) indicate the presumed depth of the electrodes based on SPW properties. (D) Density plot of the z-scored SPW peak amplitude and polarity during rest, versus normalized theta power during run in VR. Theta and SPW amplitudes were significantly correlated for both the positive polarity SPW (n=361, r = 0.24, p<10^−5^, Spearman’s rank correlation, here and subsequently, unless specified otherwise) and the negative polarity SPW (n=737, r = −0.24, p<10^−10^). (E) Similar to D, but for the eta power, which shows the opposite to theta pattern (for positive SPW n=279, r = −0.53, p<10^−10^, for negative SPW, n=472, r = 0.11, p= 0.04). (F) Distributions of correlation values between the absolute value of z-scored sharp wave peak amplitude during rest and eta normalized power during run was significantly negative (−0.34±0.04, p<10^−10^, n = 70), but the same for theta was significantly positive (0.20±0.03, p<10^−5^, n = 85). Only the sessions with at least four electrodes in the hippocampus were used. Shades in C show s.e.m.

This ensemble analysis could be influenced by differences in behavior across sessions. To address this, we computed the correlation between the SPW amplitude and theta or eta power across only those electrodes that were recorded simultaneously within a session. As expected, the SPW and theta amplitudes were significantly positively correlated for the majority of sessions (Fig. 2F). But the correlation was significantly negative between SPW and eta amplitude (Fig. 2F). These results show that while theta has a lower magnitude in the CA1 cell layer and higher magnitude in the dendrites, eta amplitude shows the opposite pattern, with highest amplitude within the CA1 cell layer in VR.

These results demonstrate that eta rhythm is not the same as type 2 theta, which appears during immobility. To further confirm this, we examined the effect of the running speed on the amplitude and frequency of eta and theta (*14–16, 24, 25*). The rats’ speed profile was comparable in RW and VR (fig. S3), with a small reduction of the peak running speed in VR (*14*). Consistent with previous studies (*14–16, 24, 25*), theta amplitude (Fig. 3E) and frequency (Fig. 3F) increased with running speed in RW. Theta amplitude showed comparable increase with running speed in VR (Fig. 3E). Surprisingly, theta frequency in VR reduced significantly from immobility to low speeds and then showed no clear dependence on speed (Fig. 3F). In addition, eta amplitude was negatively correlated with running speed in RW (Fig. 3, G and I, fig. S5). In contrast, in VR, eta amplitude reduced from immobility to low speeds (Fig. 3, G and J) and then increased with running speed (Fig. 3, G and I, fig. S5). 7.5% and 43.1% of all tetrodes showed significant increase of eta amplitude with running speed in RW and VR, respectively (Fig. 3I), while this increase was comparable for theta amplitude in the RW (79.5%) and VR (81.9%). Eta frequency showed no clear relationship with the running speed in RW or VR (Fig. 3H). This suggests that type 2 theta was present during immobility in both RW and VR, which then decreased with running speeds. However, eta rhythm appeared only in VR at high speeds and its amplitude increased with running speed, thus generating a nonmonotonic speed-dependence. Thus, speed influenced theta and eta amplitude and frequency significantly differently in RW and VR (Table S1).

**Fig. 3.**
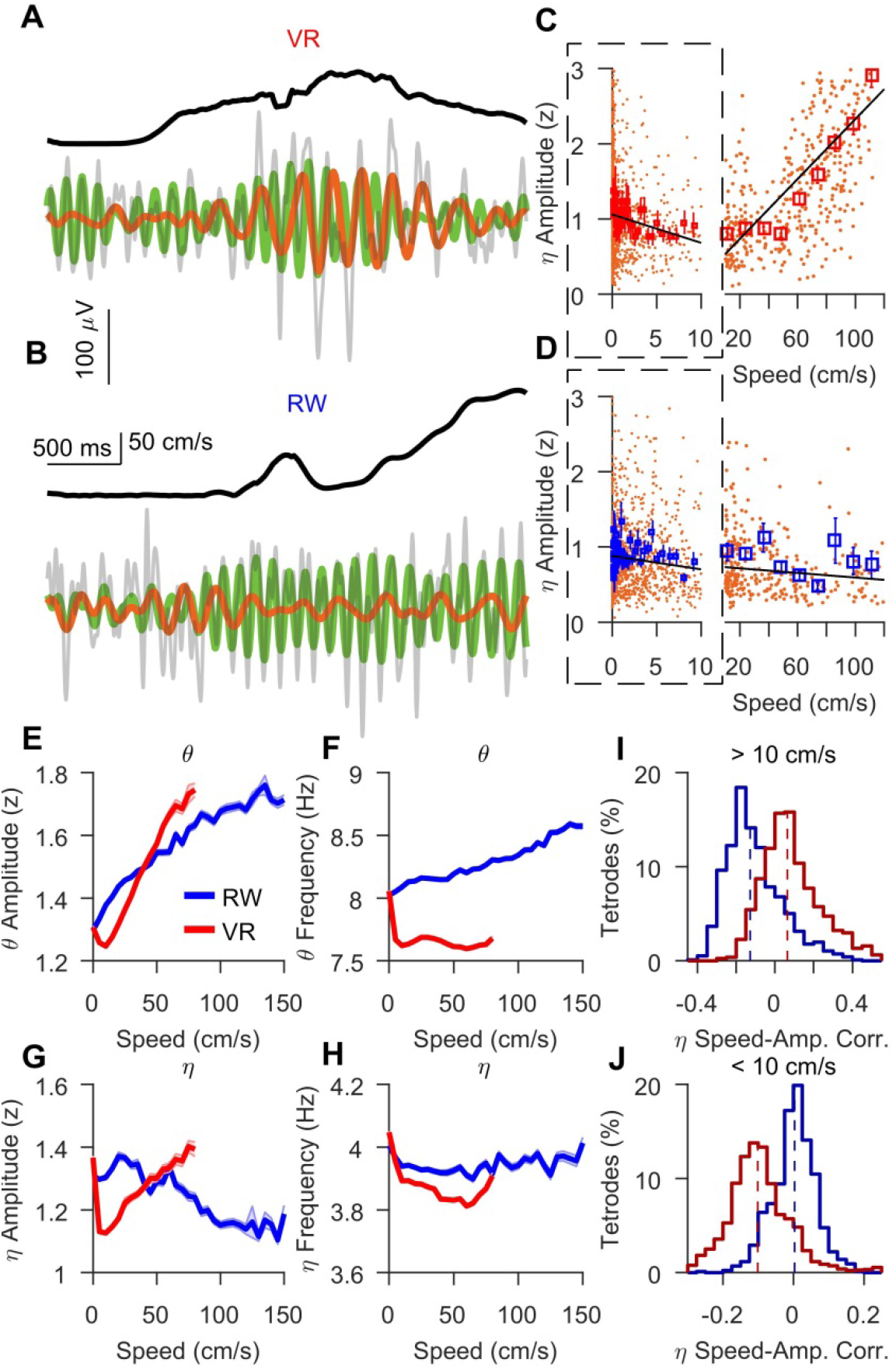
Differential effect of speed on eta amplitude and theta frequency in RW and VR. (A) Running speed of the rat (top, black) and the corresponding LFP (same format as in **Fig. 1A**) in VR. Both theta and eta amplitudes increase with speed. **(B)** Same tetrode measured in RW on the same day showing speed-dependent increase in theta, but not eta, amplitude. **(C, D)** Individual LFP eta-cycle amplitude and corresponding speed in VR (**C**) and RW (**D**) for the entire session in **A** and **B**. The broken axis separates two speed ranges – below (outlined) and above 10 cm/s. Each small dot indicates one measurement. The square dots show mean and s.e.m. in each bin in RW (blue) and VR (red). A log speed scale was used for the speed range below 10 cm/s. Linear regression fits are shown separately for both speed ranges (black lines). **(E)** Population averaged theta amplitude, showing strong increase with running speed in RW. Population averaged theta amplitude in VR first decreased at low speeds (0 vs 10 cm/s) and then increased comparable to RW (Table S1). **(F)** Same as 3E, but for theta frequency showed significant increase with running speed in RW, but in VR the frequency dropped at very low speeds (0 vs 10 cm/s), and then became speed-independent. **(G)** Same as 3E, but for eta amplitude, showing decrease in eta amplitude with increasing running speed for RW, but sharp drop in eta amplitude at low speeds (0 vs 10 cm/s), and steady increase in amplitude at higher speeds in VR. **(H)** Same as 3E, but for eta frequency, showing no clear dependence of eta frequency on running speed in both RW and VR. **(I)** Individual eta-cycle amplitudes are positively correlated with speed above 10 cm/s across tetrodes in VR (0.09 ± 0.001, p<10^−10^), but not in RW (−0.107 ± 0.005, p<10^−10^). 7.5% and 43.1% of all tetrode showed significant increase in eta amplitude with running speed in RW and VR, respectively (p<0.05). (J) Individual eta-cycle amplitudes are negatively correlated with speed below 10 cm/s across tetrodes in VR (−0.095 ± 0.003, p<10^−10^), but not in RW (0.002 ± 0.003, p = 0.1). Shaded areas and error bars denote s.e.m.

What is the dynamics of theta and eta rhythms at the level of single units? We analyzed the activity of 407 and 499 place fields obtained from the 289 RW and 459 VR recorded units, respectively (*14*). Consistent with previous studies (*14*), spikes within the place fields showed no significant difference in the degree of phase locking to the LFP theta in RW and VR (Fig. 4A). This was also true for the eta phase locking (Fig. 4B). The degree of eta phase locking was much weaker than theta – place fields that were significantly phase locked to theta (~72%) were four times more than those locked to eta (~18%) in both RW and VR (Fig. 4, C and D). The preferred theta phase was also similar in RW and VR, with most of the place fields preferentially spiking near theta peak (*4, 26*) (Fig. 4, E, F).

**Fig. 4.**
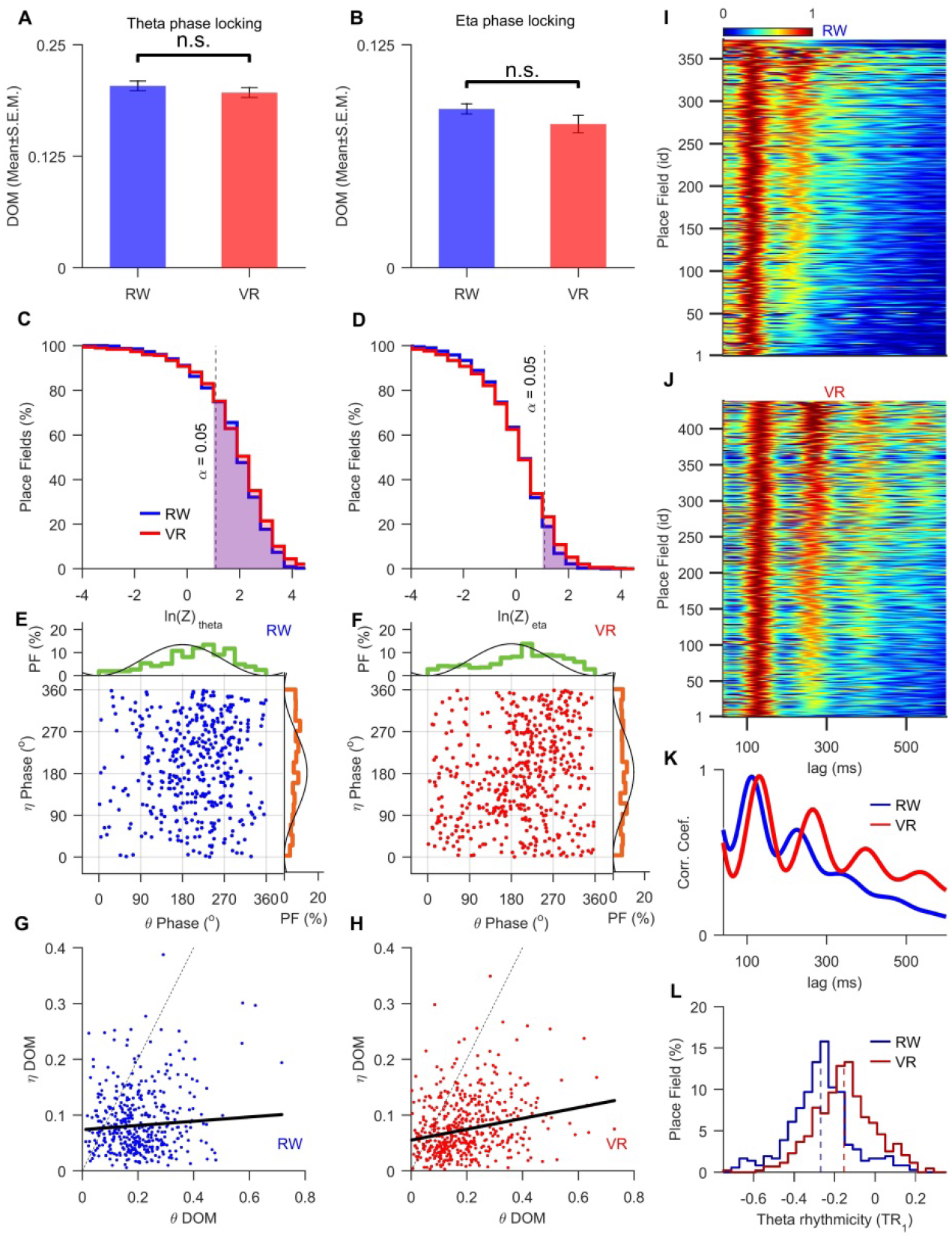
Enhanced theta rhythmicity but not eta modulation of CA1 place cells in VR. (A) Magnitude of theta phase locking in RW (0.2±0.005, n = 407) and VR (0.196±0.006, n = 499) are not significantly different (p=0.07, χ^2^ = 3.26). (B) Same as 4A, but for eta phase locking in RW (0.089±0.003) and VR (0.08±0.002) also showing no significant difference (p=0.076, χ^2^ = 4.93). (C) Cumulative distribution of log-transformed Rayleigh’s Z computed for theta modulation of the place cells. This was significant (shaded area) at 0.05 level for a majority of cells in RW (n = 297, 72.9%) and VR (n = 360, 72.1%). (D) Same as 4C, but for eta band in RW (n = 60, 14.7%) and VR (n = 100, 20.04%). (E) Relationship between preferred theta and eta phases of place fields in RW (n = 407, r = 0.16, p = 0.0014, circ-circ corr.). The corresponding distributions (circ. (mean± std.)) of preferred theta (223.65±1.40°, MVL = 0.50, green) and eta (241.53±1.39°, MVL = 0.40, brown) phases of place fields in RW are shown. A reference theta/eta cycle is plotted in black (LFP positive polarity is downward). (F) Same as in 4E, but for VR (n = 499, r = 0.15, p < 0.001). The corresponding distributions of preferred theta (231.71±1.32°, MVL = 0.28, green) and eta (166.34±1.30°, MVL = 0.06, brown) phases in VR are shown. The preferred distributions are significantly different for theta (p = 0.02, V = 0.132, Kuiper’s test, 5) and eta (p = 0.01, V = 0.175, Kuiper’s test) between RW and VR. (G) No significant correlation was found between theta and eta depth of modulation of spikes (DoMs), defined as the magnitude of phase locking (see methods), within the place fields in RW (n = 407, r = −0.025, p= 0.605, partial Spearman correlation factoring out number of spikes). (H) Similar to 4G, but for VR showed significant positive correlation between eta and theta DoMs (n = 499, r = 0.19, p<10^−5^). (I) Autocorrelograms of spike trains (corrected by the overall ACG decay, see methods) ordered according to the increasing TR1 values for the place fields in RW. The autocorrelograms are normalized by the amplitude of their theta peak to allow easy comparison. (J) Same as 5I, but for VR, showing more theta peaks, i.e. greater rhythmicity, than in RW. (K) Population average of autocorrelations shows greater theta rhythmicity in VR than in RW. (L) Distribution of the theta rhythmicity (TR1) index was significantly greater (p<10^−10^, *X*^2^ = 123.3) in VR (−0.151±0.07) than RW (−0.275±0.158).

In contrast to theta, place fields preferred a wide range of eta phases in both RW and VR (Fig. 4, E and F). The circular correlations between place fields’ eta and theta phase preferences were equally weak in RW (Fig. 4E) and VR (Fig. 4F). The LFP theta and eta phase locking of place fields were uncorrelated in RW (Fig. 4G). In VR, they were significantly but weakly correlated (Fig. 4H). The co-modulation of theta and eta in VR suggests that the two rhythms maybe coupled in VR but not in RW. Consistently, LFPs showed significant phase-phase coupling between eta and theta in VR but not in RW (fig. S6).

To capture the degree of sustained rhythmicity, we computed the autocorrelograms (ACG) of the spike times within the place fields. This revealed greater rhythmicity in VR, exhibiting several peaks at theta and its harmonics, compared to RW (Fig. 4, I, J and K, fig. S7). To quantify this, we computed the theta rhythmicity index (TR) defined as the differences between adjacent peak amplitudes in ACG, normalized by the greater of both. This was done for up to the fourth ACG peak (TR_1_, TR_2_, TR_3_, fig. S8). The rhythmicity index computed for the first ACG peak (TR_1_) is related to the theta skipping index (*27, 28*) and was (80%) greater in VR than in RW (Fig. 4L).

The rhythmicity index could be influenced by nonspecific properties of place fields, such as spatial width. To overcome this, we did additional analyses by fitting a Gaussian mixture model to the ACG to obtain an unbiased estimate of rhythmicity (see methods, fig. S7). The model-corrected rhythmicity indices (TR_1_, TR_2_ and TR_3_) were consistently larger in VR than in RW (Fig. 4L, fig. S8), revealing sustained increase in rhythmicity in VR. As an additional confirmation, we factored out the contribution of place fields’ widths and the results remain qualitatively unchanged (fig. S9). The enhanced theta rhythmicity in VR was related to the magnitude of the eta and theta phase locking (fig. S10).

Theta is strongly influenced by inputs from the medial septum (*6, 25, 29*), which targets hippocampal inhibitory neurons. Hence, we examined the rhythmicity of 34 and 174 putative inhibitory interneurons in RW and VR, respectively. The magnitudes of both theta (Fig. 5A) and eta (Fig. 5B) phase locking of the interneurons were nearly twice as large in VR as compared to RW. While all interneurons showed significant theta phase locking in both RW and VR (Fig. 5C), the significantly eta phase locked interneurons in VR (66.6%) were far greater than in RW (35.3%) (Fig. 5D), which is unlike the pyramidal cells (Fig 4D). Furthermore, theta phase preference of the population of interneurons was similar in both worlds (Fig 5, E and F). But, the population of interneurons, and not pyramidal neurons, showed greater eta phase preference in VR by preferentially firing near the eta peak (Fig. 5, E and F). As a result, a strong positive circular correlation was seen between eta and theta phase preferences of interneurons in VR (Fig. 5F) but not in RW (Fig. 5E). Similarly, eta to theta co-modulation of interneurons was stronger in VR (Fig. 5H) than in RW (Fig. 5G) and far greater than in pyramidal neurons. In addition, the interneurons’ ACG in VR showed greater rhythmicity than in RW (Fig. 5, I, J and K), and significantly greater TR (Fig. 5L, figs. S8 and S11). Interneurons with higher theta rhythmicity showed increasingly more theta and eta phase locking in VR (fig. S10), but not in RW.

**Fig. 5.**
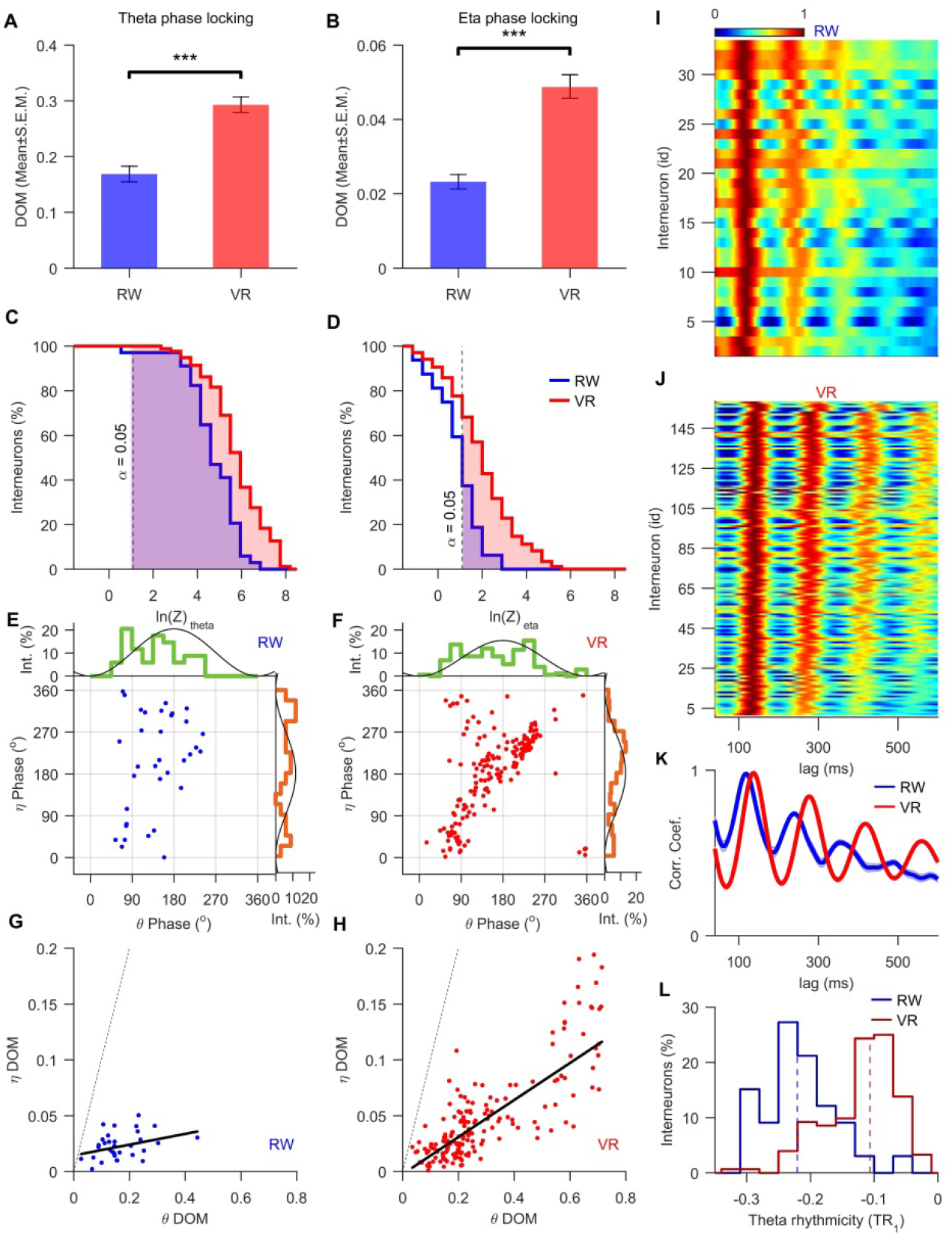
Enhanced theta rhythmicity, eta and theta modulation of interneurons in VR. (A) Magnitude of the theta phase locking of interneurons in VR (0.29±0.014) is significantly greater than in RW (0.16±0.014) (χ^2^ = 16.35, p<0.001) by 81%. (B) Same as 5A, but for eta, showing significantly greater (150%) phase locking of interneurons in VR (0.05±0.003) compared to RW (0.02±0.001) (χ^2^ = 14.25, p<0.001). (C) Cumulative distribution of the log-transformed Rayleigh’s Z of theta modulation for interneurons shows significantly modulated (shaded area) cells in VR (n = 174, 100%) and in RW (n = 33, 97.05%) at p< 0.05 (dashed line). (D) Same as 5C, but for eta band shows 31.37% more cells are significantly modulated in VR (n = 116, 66.66%) than in RW (n = 12, 35.29%). (E) Relationship between preferred theta and eta phases of interneurons in RW (n = 34, r = −0.44, p = 0.011, circ-circ corr.). The corresponding distributions (circ. (mean± std.)) of preferred theta (137.82±1.32°, MVL = 0.48, green) and eta (291.67±1.39°, MVL = 0.21, brown) phases are specified. A reference theta/eta cycle is plotted in black (LFP positive polarity is downward). (F) Same as in 5E, but for VR (n = 174, r = 0.35, p = 1.8×10^^−6^). The corresponding distributions of preferred theta (155.4±1.38°, MVL = 0.42, green) and eta (219.41±1.35°, MVL = 0.29, brown) phases in VR are shown. The distributions are significantly different for theta (p = 0.02, V = 0.132, Kuiper’s test) and eta (p = 0.05, V = 0.339, Kuiper’s test) preferred phases between RW and VR. (G) No significant correlation between eta and theta DoMs of interneurons was seen in RW (n = 34, r = 0.28, p = 0.11, partial correlation factoring out number of spikes). (H) Same as 5G, but in VR showed strong positive correlation (n = 174, r = 0.75, p < 10^−10^, partial correlation factoring out number of spikes). I, Corrected ACG ordered according to the increasing TR_1_ values for RW. The autocorrelograms are normalized by their first theta peak values as for the place cells. (J) The same as 5I, but for VR, showing more theta peaks, i.e. greater rhythmicity, than in RW. (K) Population average of autocorrelations showed greater theta rhythmicity in VR compared to RW. (L) Histograms of the TR_1_ distributions in VR (−0.118±0.004) is 82% greater (p<10^−10^, *X*^2^ = 47.31) than in RW (− 0.215±0.009).

These results revealed the crucial role of multisensory inputs in hippocampal rhythmogenesis. We found that rodents’ dorsal hippocampal CA1 can simultaneously exhibit two distinct slow oscillations, eta and theta, when rats are running in a visual VR, which is different from RW where the hippocampus shows only theta. The eta rhythm in rodent VR may be related to the irregular bouts of 1-5 Hz oscillations reported in humans and nonhuman primates while they are immobile, and performing hippocampus-dependent tasks in VR (*10, 11, 30–32*). On the other hand, eta amplitude in our studies is greater during locomotion than immobility. When humans walk, a higher, ~8 Hz theta oscillation appears in hippocampal LFP, which is either absent or substantially reduced in VR (*30, 33*). In contrast, we found that the ~8 Hz theta is present not just in RW but in VR as well, and theta rhythmicity is even greater in VR than in RW.

These differences between humans and rodents could arise due to several factors. One possible reason is that humans were immobile in these VR studies and could make only restricted eye and hand movements, compared to rats in our studies, which ran similarly in RW and VR making the full sets of running movements. However, because of the body-fixed condition in VR, the linear acceleration is minimized. Hence, we hypothesize that running movements of the body, without significant linear acceleration, is sufficient to induce or enhance eta and theta rhythms. In particular, amplitude of the eta rhythm is enhanced by running movement in VR, which is different from the type 2 theta occurring during immobility, whose amplitude reduces during running in RW. Theta amplitude and rhythmicity is also enhanced by running in VR. We hypothesize that the inclusion of linear acceleration in RW abolishes eta rhythm and reduces theta rhythmicity, in part, via interference of different sources of slow oscillations. This is consistent with studies showing that several brain areas are involved in theta rhythm (*9*).

The eta rhythm is unlikely to be the ~4 Hz synchrony between hippocampus and prefrontal cortex found in studies conducted in RW (*34*), where eta is not seen in hippocampal LFPs. The eta rhythm is also unlikely to be related to the respiration related rhythm (*35*), since the respiratory rhythm is weaker in the cell layer and stronger below the cell layer which was not found for eta rhythm. This further demonstrates that eta is not volume conducted signal from other brain areas. In fact, eta was highest in the CA1 cell layer and lower above and below, in the dendrite rich regions. This is in stark contrast to theta, which shows the opposite of soma-dendrite amplitude profile.

We hypothesize that the eta rhythm is likely generated within the CA1 cell layer by a local network of excitatory-inhibitory neurons. Accordingly, it is not the pyramidal neurons but the inhibitory interneurons’ activity that was differentially modulated by eta in VR compared to RW. This is further supported by several *in vivo* and *in vitro* studies demonstrating the role of CA1 interneurons in hippocampal slow oscillations (*36–40*). We hypothesize that the network oscillations of CA1 excitatory-inhibitory network in VR slows down due to the shutdown of a large number of pyramidal neurons in VR (*14*). This can also explain the slower theta frequencies and the emergence of locally generated eta rhythm in VR. Coupled with theta, eta can enhance the rhythmicity and alter the speed dependence of the theta rhythm in VR. This mechanism can be most prominent at the large dendritic branches of the pyramidal cells, where theta is largest. Recent theories suggest that memories are encoded on segments of dendritic branches in pyramidal neurons flanked by inhibitory synapses (*41, 42*). This would result in decoupling of the dendritic activity from the soma, as observed recently (*43*). The interaction between multisensory codes across different dendritic branches of pyramidal neurons would result in reduced theta rhythmicity and eta modulation in the RW.

The presence of the slow oscillation, nearly half as slow as theta, could segregate activity of hippocampal neural populations into parallel streams of hippocampal information processing throughout theta cycles (*27, 28, 44*). Moreover, hippocampal eta activity may facilitate long-range inter-structural coordination, since the 4 Hz rhythm is the dominant activity mode in cortical structures (*34, 45*). The presence of eta rhythm and enhanced theta rhythmicity in VR would influence neural synchrony and via NMDAR-dependent synaptic plasticity (*46, 47*) in dendritic branch specific fashion (*43*), to alter hippocampal circuit and learning (*42*). Thus, these findings elucidate the role of multisensory inputs in governing hippocampal slow oscillations and reveal the surprising enhancement of hippocampal theta rhythmicity and eta modulation in virtual reality.

## Acknowledgments

We thank Pascal Ravassard, Ashley Kees, Lavanya Acharya and Ayaka Hachisuka for providing the experimental data; Zahra Aghajan and Bernard Willers for help with the analyses; Shonali Dhingra, Krishna Choudhary, current and former Mehta lab members for discussions, careful reading of the manuscript and valuable comments.

## Author contributions

K.S. and M.R.M. designed the study, M.R.M. advised on all aspects of the analysis and experiments. K.S. analyzed the data, generated the figures. K.S. and M.R.M. wrote the paper. Competing interests: Authors declare no competing interests.

## Data and materials availability

All data is available in the main text or the supplementary materials upon reasonable request.

## Supplementary Materials

### Materials and Methods

#### Subjects and surgery

Detailed methods have been described previously(*14*). Briefly, seven, adult, male, Long-Evans rats (approximately 3.5 months old at the start of training) were implanted with 25-30 g custom-built hyperdrives containing up to 22 independently adjustable tetrodes (13 μm nichrome wires) positioned over both dorsal CA1 areas (−4.0 mm A.P., 2.4 mm M.L. relative to bregma). Surgery was performed under isoflurane. Analgesia was achieved by using Lidocaine (0.5mg/kg, sc) and Buprenorphine (0.03mg/kg, ip). Dura mater was removed and the hyperdrive was lowered until the cannulae were 100 μm above the surface of the neocortex. The implant was anchored to the skull with 7-9 skull screws and dental cement. The occipital skull screw was used as ground for electrophysiology. Electrodes were adjusted each day until stable single units were obtained. Positioning of electrodes in CA1 was confirmed through the presence of SPW ripples during immobility.

#### Virtual reality and real world tasks

The virtual environment consisted of a 220×10 cm linear track floating 1 m above the virtual floor and centered in a 3×3×3 m room(*13, 14*). Alternating 5 cm-wide green and blue stripes on the surface of the track provided optic flow. A 30×30 cm white grid on the black floor provided parallax-based depth perception. Distinct distal visual cues covered all 4 walls and provided the only spatially informative stimuli in the VR. In RW, rats ran back and forth on a 220×6 cm linear track that was placed 80 cm above the floor. The track was surrounded by four 3×33 m curtains that extended from floor to ceiling. The same stimuli on the walls in the virtual room were printed on the curtains, thus, the distal visual cues were similar in RW and VR.

#### Data acquisition, LFP processing, spike detection, sorting and cell classification

Spike and LFP data were collected by 22 independently adjustable tetrodes. Signals from each tetrode were digitized at 32 kHz and wide band pass-filtered between 0.1 Hz and 9 kHz (DigiLynX System, Neuralynx, MT). This was down-sampled to 1.25 kHz to obtain the LFPs, or filtered between 600-6000Hz for spike detection. LFP positive polarity was downward(*14*). Unless otherwise stated, the bandpass LFP filtering was done by using a zero-lag forth order Butterworth filter. Spikes were detected offline using a nonlinear energy operator threshold(*14*). After detection, spike waveforms were extracted, up-sampled fourfold using cubic spline, aligned to their peaks and down-sampled back to 32 data points. PyClust software was used to perform spike sorting(*48*). These were then classified into putative pyramidal neurons and interneurons based on spike waveforms, complex spike index and rates(*14*).

Offline analyses were performed using custom code written in MATLAB (MathWorks).

#### Analysis of LFP and spike data during behavior

Running epochs were defined as continuous periods of running (>10 cm/s) for 2 sec. or more. Immobility was defined as periods of low speed (<2.5 cm/s) for 2 sec. or more. Hence, the low speed range, which excluds periods of immobility, was taken as a range from 5 to 15 cm/sec. In addition, correlation coefficients of the amplitudes of the different frequency bands with speeds were computed below and above 10 cm/sec to capture dynamics during transition periods from the rest to run and running epochs.

Spectral analysis of oscillatory activity was computed using a multitaper method(*49, 50*) by Chronux toolbox (http://chronux.org). The window size of 4 sec. (average running time during the task) and 3–5 tapers were used with a 75% overlap over frequencies ranging from 0.5 to 30 Hz. The spectral power was computed separately during running epochs and immobility states. The power spectral index was computed as the difference of power between running epochs and immobility at each frequency over their sum (Fig. 1H, 2B, Fig. S1). Significance of the power difference between running epochs and immobility in the theta and eta bands was determined using Kruskal-Wallis nonparametric test (α = 0.01). To reduce nonspecific effects, the power spectra of each tetrode were normalized by average power in 0.5-30 Hz range on that tetrode separately for the running epochs and immobility states. Theta and eta power peaks were detected using peak prominence of 0.01 or more within the respective frequency bands (findpeaks.m from signal toolbox in MATLAB). The prominence was defined as the height of the peaks at the levels of highest troughs (Mathworks). With few exceptions this led to the detection of eta peaks predominantly during running epochs in VR. The prominence of eta index peaks greater than 5 percentile of the theta index peaks was considered as significant. Peak power was computed as an average power within 1 Hz at the detected peak.

This power spectrum based method requires comparatively long periods of unitary behavior (e.g. run or stop) over which the power spectra are computed. To obtain an estimate of the instantaneous values of eta and theta bands, we filtered the LFP data in either eta (2.5-5.5 Hz) or theta (6-10 Hz) ranges and computed its Hilbert transform. Amplitude difference index was computed as the difference of the mean amplitude in theta (or eta) band during high (30-60 cm/s) and low (5-15 cm/s) speed runs, divided by their sum (Fig. 1 c, d). Significance level of theta (or eta) modulation of LFP was determined by comparing the distributions of the LFP amplitude in theta (or eta) band during high (30-60 cm/s) versus low (5-15 cm/s) speed runs, and using a nonparametric Kruskal-Wallis test. Alternative, non-parametric estimate was also done by computing robust regression fits between amplitude envelope and speed.

Theta frequency was computed using three methods: cycle detection using Hilbert transformed phase jumps, the derivative of Hilbert transform phase, and the short time Fourier transform. The cycle method results are reported in the main text (Fig. 3, Extended Data fig. 3).

#### Sharp wave and ripple detection

To estimate the electrode depth we did SPW ripple analysis during periods of immobility in baseline sessions preceding the tasks. LFP data was filtered in ripple (80-250 Hz) band. This signal was z-scored by subtracting the mean value and dividing by the standard deviation of the ripple band LFP. Double-threshold crossing method was applied to the ripple band LFP(*51, 52*) to quantify ripple events. All time points with ripple amplitude larger than a first threshold (mean ripple amplitude + 3 standard deviations) were identified as part of a ripple event. Only events with a peak value larger than a second threshold (mean + 10 standard deviations) and duration larger than 30 ms were retained (Fig. 2c). Ripple events separated by less than 50ms were stitched together. To determine the concomitant SPW, LFP were filtered in 6-25 Hz range. These signals were z-scored and its amplitude detected at the times of each associated ripple peak power. These SPW were averaged across all the ripples in a session to obtain the average SPW (Fig. 2c).

#### Place field detection

A unit was considered track (goal) active if its mean firing rate on track (goal) was at least 1 Hz. Opposite directions of the track were treated as independent and linearized. A place field was defined as a region where the firing rate exceeded 5 Hz for at least 5 cm. The boundaries of a place field were defined as the point where the firing rate first drops below 5% of the peak rate (within the place field) for at least 5 cm, and exhibits significant activity on at least five trials(*14*).

#### Phase locking detection and characterization

Instantaneous amplitudes and phases were estimated by Hilbert transform of the filtered signals as below:

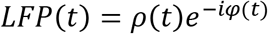

where *ρ(t)* is instantaneous amplitude and *φ(t)* is instantaneous phase.

Rayleigh circular uniformity test was computed to test significance of phase locking. The first circular moment was given as: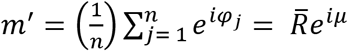, where *φ*_*j*_ are phases of *n* spikes. The preferred LFP phase of spikes is thus given by *μ* and the magnitude of phase locking was given by 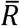. The magnitude of phase locking was defined as depth of modulation (DoM). The Rayleigh statistics 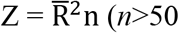, only neurons with at least 50 spikes were used), and the probability of the null hypothesis of sample uniformity (p = e^−Z^) was applied(*53,54*).

#### Spike autocorrelogram (ACG), Gaussian mixture model (GMM) fit and theta rhythmicity index (TR) calculation

Spike-time autocorrelograms were computed using accuracy of 1ms, smoothed by 20 ms Gaussian function, and normalized by the number of spikes to obtain probability at lags. Autocorrelograms *Y(t)* were fit using the following Gaussian mixture model(*55–57*):

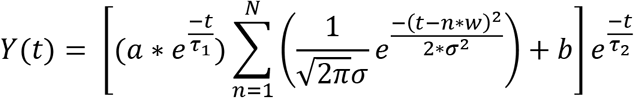

Where *t* is the autocorrelation lag time (ranging from 60-600 ms) and *a*, *τ*_1_, *w*, *σ*, *b*, *τ*_2_ are the fit parameters. The Gaussian terms are used to fit theta peak and its harmonics (*w* is a first theta lag). *τ*_1_ and *τ*_2_ are the exponential decay constants for the magnitude of rhythmicity and overall ACG falloff rate due to finite amount of data, respectively. *a* is the rhythmicity factor and *b* is constant background or non-rhythmic component. *N*=5 five Gaussians *n* were used to fit the ACG. The amplitude of the first Gaussian (*n=1*) provides an estimate of theta modulation, while removing nonspecific effects arising from the duration of the place field and the duration of recording. Theta rhythmicity was defined as TR(n) = (amplitude(n+1) – amplitude (n))/max(amplitude (n), amplitude (n+1)) where *n* is a ACG peak amplitude at theta or its harmonics and varies from *1* to *3*. Theta skipping (*27, 28*) was defined as TR with *n*=1.

#### Statistics

Unless otherwise stated, statistical significance between two distributions of linear variables was evaluated using nonparametric Kruskal-Wallis test. Tests for populations significantly different from zero were also performed using the nonparametric Kruskal-Wallis test. Average values are reported in the form mean ± s.e.m. unless otherwise stated. Median values of histograms are depicted as a dashed line in all main figures. Circular statistics were computed using the CircStat toolbox. Binomial confidence interval was computed using the Clopper-Pearson method from statistics toolbox in MATLAB (binofit.m). To reduce the contribution of outliers, unless otherwise stated we used nonparametric Spearman’s rank correlation to compute all correlation coefficients including partial correlations.

## Supplementary figures

**Fig. S1.**
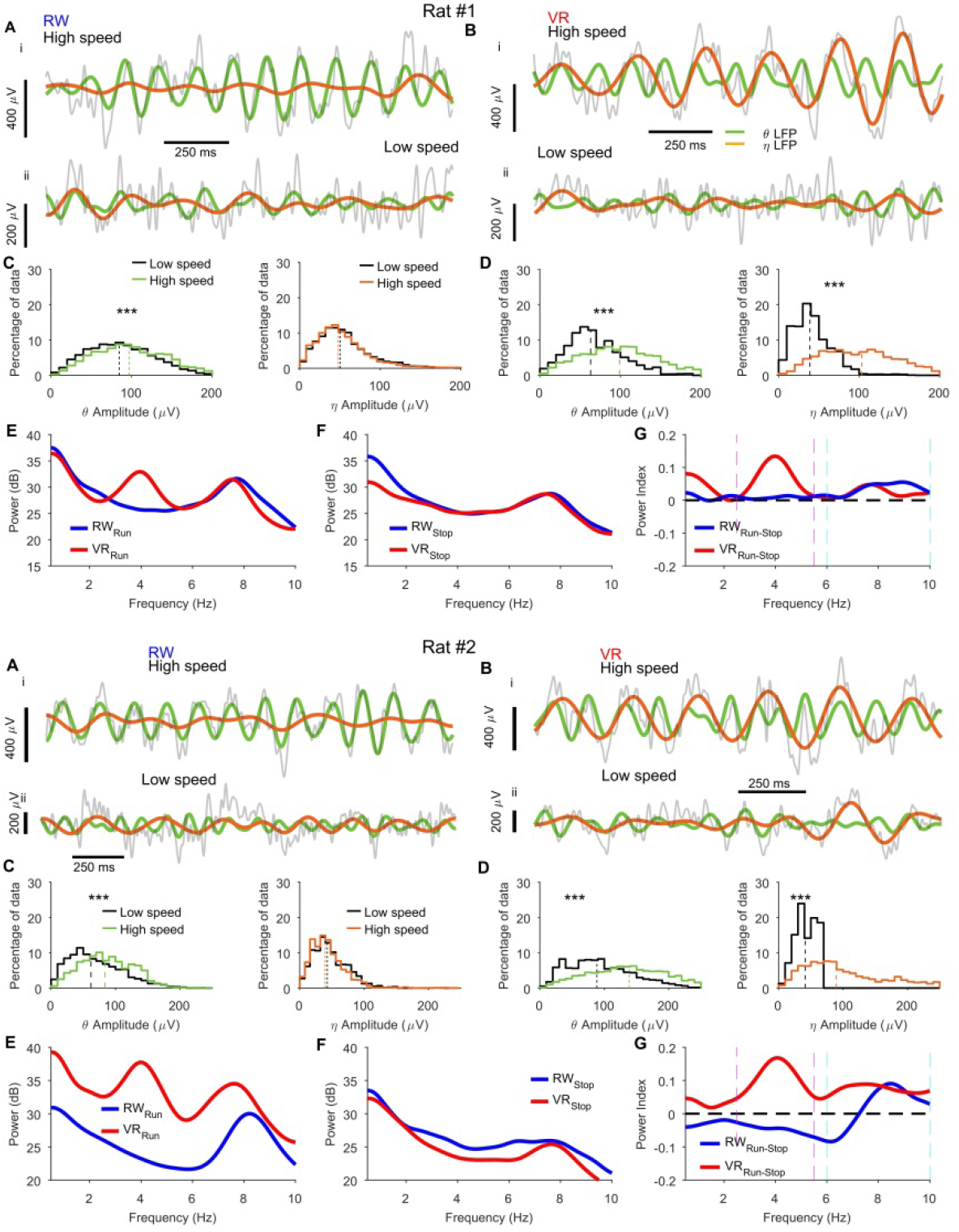
Additional examples of ~4 Hz eta oscillation during running in VR, but not in RW. The data were recorded from rat #1 and rat #2. Similar format as Fig. 1. (**A, B)** Traces of LFP, raw (grey), filtered in theta (6-10 Hz, green) and eta (2.5-5.5 Hz, brown) bands during high-speed (above 15 cm/s) running on track (top, i) and at low-speeds (below 15 cm/s) (bottom, ii) recorded on the same tetrodes in the same day RW (**A**) and VR (**B**). (**C**) Amplitude envelope distribution during high-(30-60 cm/s) and low-(5-15 cm/s) speed runs for the theta (left panel) and eta (right panel) bands in RW. Theta amplitude was significantly (rat #1, p < 10^−10^, *X*^2^ = 822.14; rat #2, p < 10^−10^, *X*^2^ = 218.0) larger at high speeds than low speeds, whereas eta amplitude was slightly smaller at high speeds (rat #1, p = 10^−10^, *X*^2^ = 49.5; rat #2, p <10^−10^, *X*^2^ = 359.5). (**D**) Similar to **C**, but for VR showing large and significant increase in both eta (rat #1, p < 10^−10^, *X*^2^ = 7942.7; rat #2, p < 10^−10^, *X*^2^ = 279.76) and theta (rat #1, p < 10^−10^, *X*^2^ = 5542.9; rat #2, p < 10^−10^, *X*^2^ = 259.14) amplitudes at higher speeds. (**E, F**) Power spectra of the example LFPs in RW (blue) and VR (red) during running (**E**) and immobility (**F**). (**G**) Power index, during run compared to stop, showing prominent peaks in both eta and theta bands in VR (red) and only in theta band in RW (blue). (*** p < 10^−10^).

**Fig. S2.**
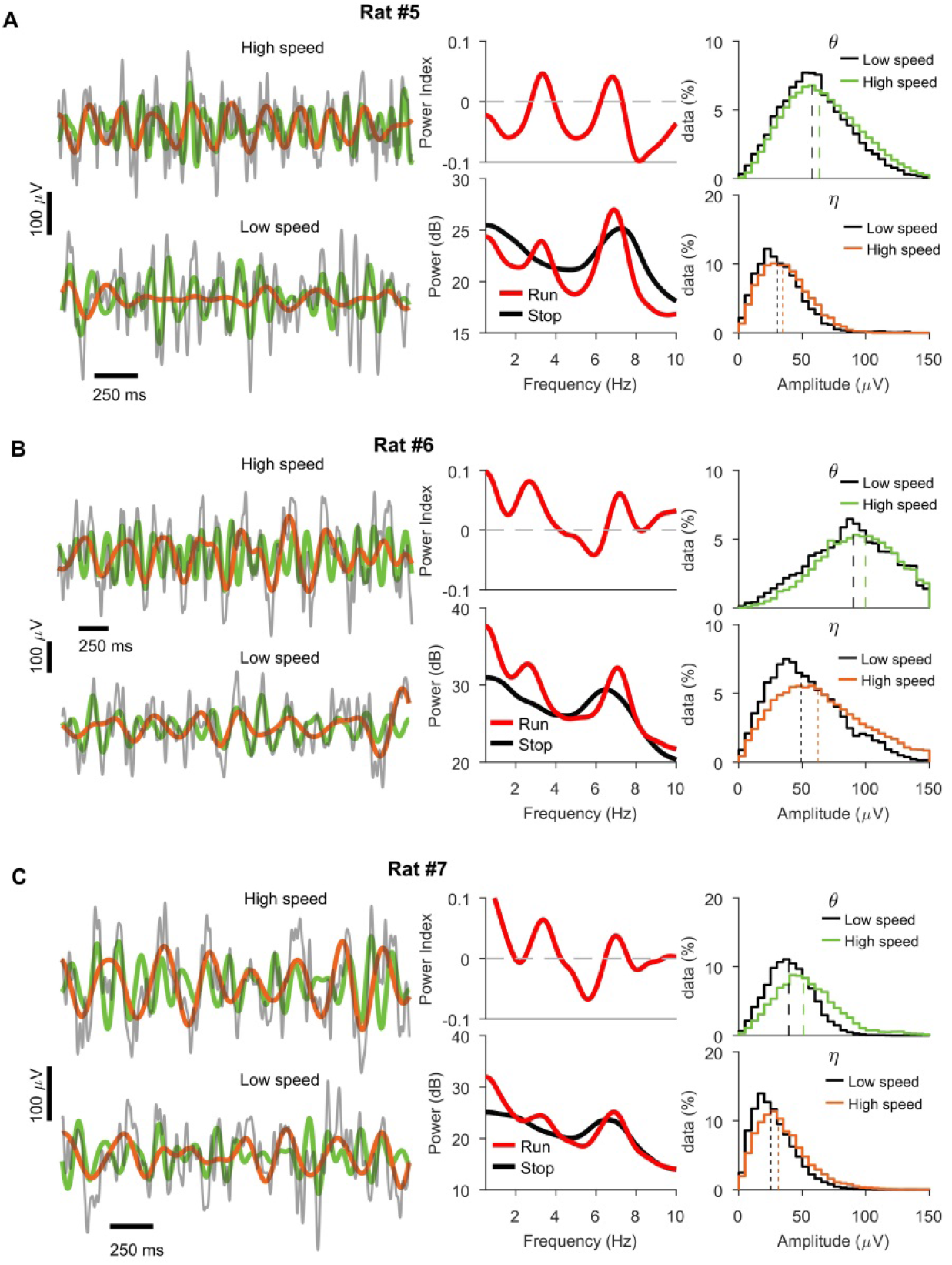
Additional examples of ~4 Hz eta oscillation during running in VR. The data were recorded from rat #5, rat #6 and rat #7. Similar format as Fig. 1. (**A, B, C) Left**, Traces of LFP, raw (grey), filtered in theta (6-10 Hz, green) and eta (2.5-5.5 Hz, brown) bands during high-speed (above 15 cm/s) running on track (top) and at low-speeds (below 15 cm/s) (bottom) recorded in the VR. **Middle**, (bottom panel) Power spectra of the example LFPs in VR during running (red) and immobility (black), and (top panel) power index, during run compared to stop, showing prominent peaks in both eta and theta bands in VR (red). **Right**, Amplitude envelope distribution during high-(30-60 cm/s) and low-(5-15 cm/s) speed runs for the theta (top panel) and eta (bottom panel) bands in VR. Theta amplitude was significantly (rat #5, p <10^−10^, *X*^2^ = 1430.5; rat #6, p <10^−10^, *X*^2^ = 4357.1; rat #7, p <10^−10^, *X*^2^ = 2661.0) larger at high speeds than low speeds, whereas eta amplitude was slightly smaller at high speeds (rat #5, p < 10^−10^, *X*^2^ = 3434.0; rat #6, p <10^−10^, *X*^2^ = 12250.0; rat #7, p <10^−10^, *X*^2^ = 1997.3).

**Fig. S3.**
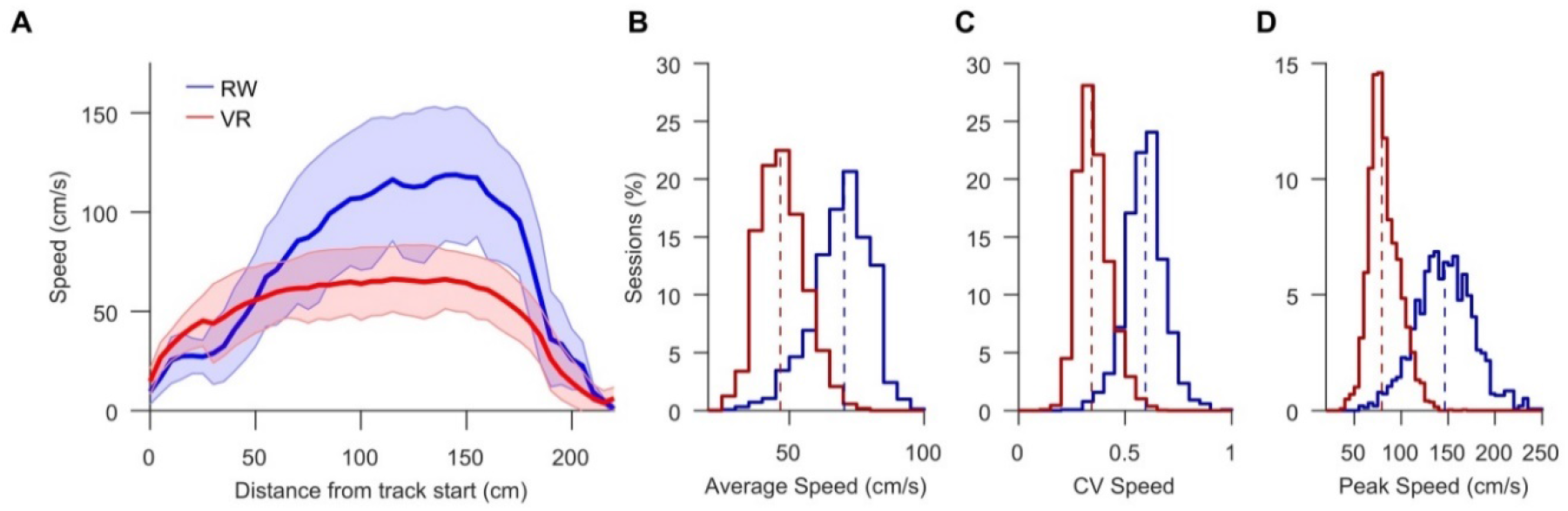
Running speed in the linear track in RW and VR. **(A)** Running speed (means ± SD) of the rats as a function of position on a 2.2-m-long linear track for RW (blue) and VR (red). Although the rats were faster in RW, their behavior was similar, reliably reducing speed before reaching the end of the track (n = 49 sessions in RW, n = 121 sessions in VR). (**B**) Average speeds in RW (69.42±0.27) were significantly greater (p <10^−10^, χ2 = 2502.3) than in VR (47.00±0.15). (**C**) CV of running speeds in RW (0.59±0.0023) were significantly greater (p <10^−10^, χ2 = 2933.6) than in VR (0.35±0.0012). (**D**) Average speeds in RW (147.13±0.02) were significantly greater (p <10^−10^, χ2 = 2872.7) than in VR (81.48±0.0046).

**Fig. S4.**
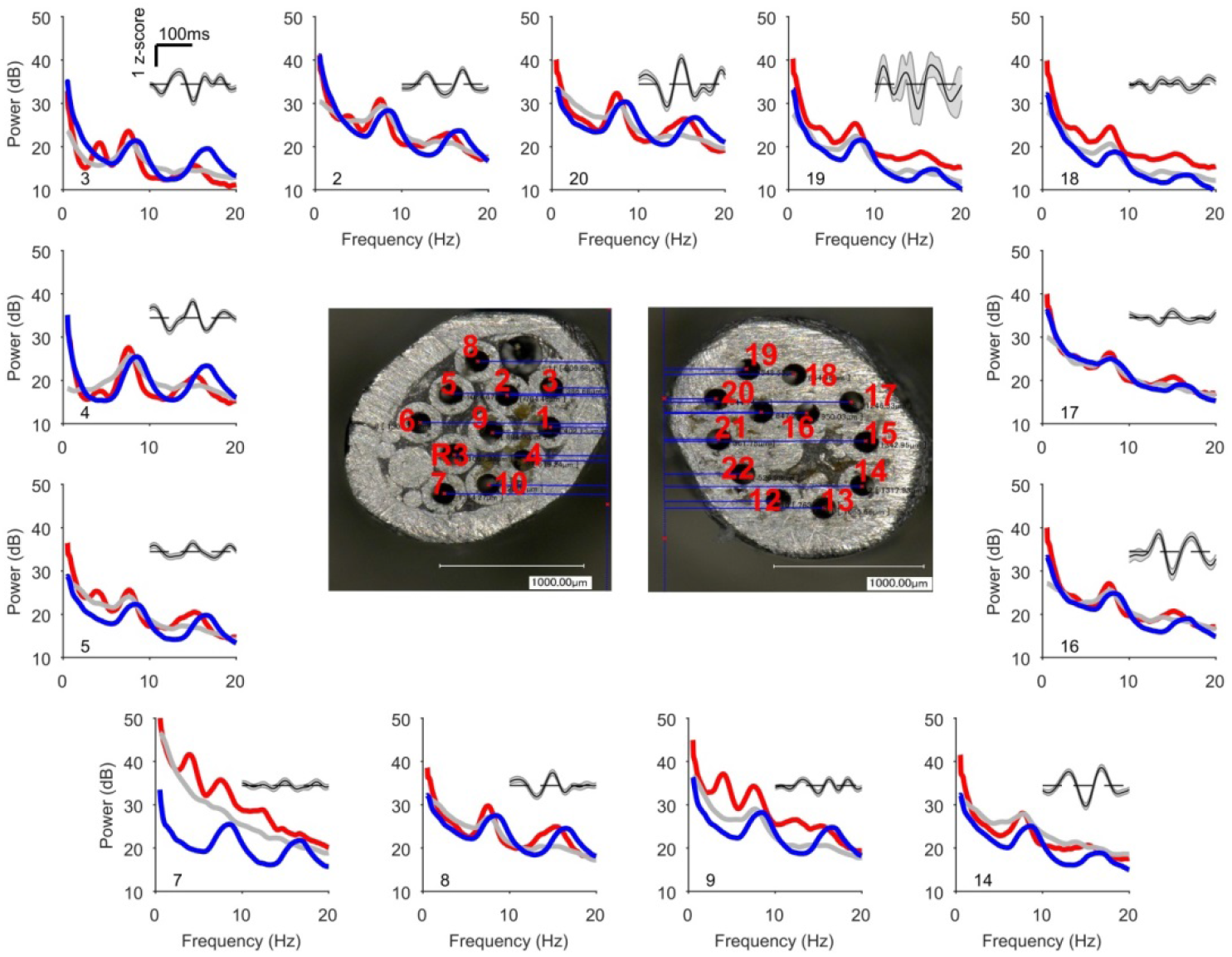
Prominent eta band peak appears only during running in VR on tetrodes with small SPW, independent of the planer position of the electrodes. LFP power spectra for simultaneously recorded tetrodes are shown during running (red) and immobility (grey) in VR. Power spectra of the same tetrodes during running in RW are also shown (blue). Average z-scored sharp-waves computed from the baseline session preceding the VR session are shown for each tetrode (grey inset). Tetrode numbers are shown at left bottom corner of the power spectra. **Center**: Pictures of the bilateral cannulae with tetrode numbers.

**Fig. S5.**
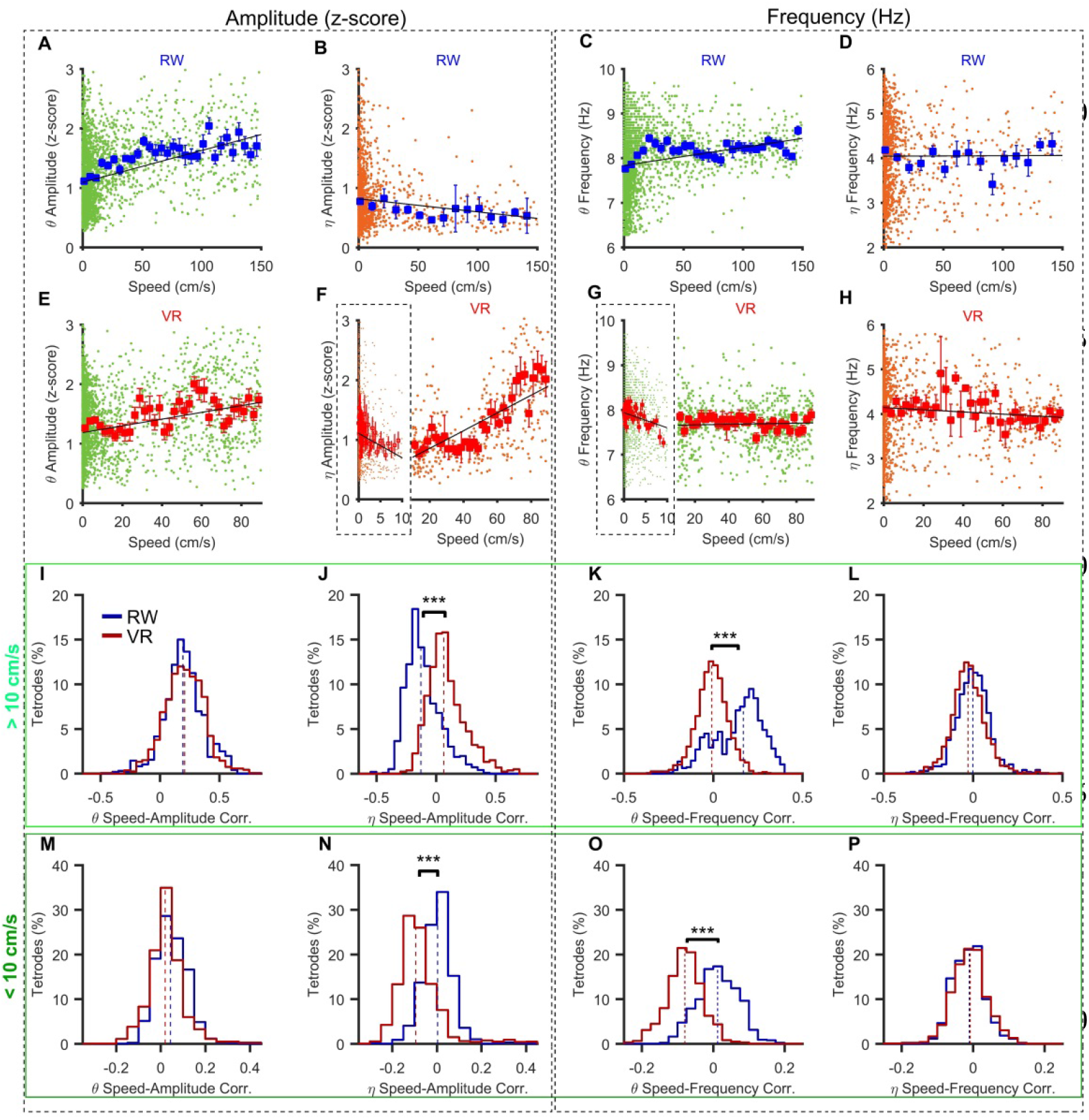
Speed dependence of theta and eta amplitude and frequency. Two more example data, similar format as Fig. 3. (**A-H**) LFP theta and eta amplitude (left two columns) and frequency (right two columns) as a function of running speed. Individual theta- and eta-cycle amplitudes (**A, B, E, F**) and frequencies (**C, D, G, H**) from a single LFP in the same day RW (**A-D**) and VR (**E-H**) recordings for an example electrode are shown as a function of speed. **(A, E)** LFP theta-cycle (green) amplitudes and corresponding speeds in RW (**A**) and VR (**E**). (**B**, **F**) Speed modulation of eta-cycle (brown) amplitudes in RW (**B**) and VR (**F**). (**C, G)** Speed modulation of theta-cycle frequency in RW (**C**) and VR (**G**). (**D, H)** LFP eta-cycle frequency speed modulation in RW (**D**) and VR (**H**). Mean and s.e.m. are shown for both RW (blue) and VR (red). Because of a near exponential distribution of the amount of data as a function of low speeds, a log speed scale was used for low speeds (outlined parts in (**F, G**)). Linear regression fits are plotted (black lines). (**F, G**) The broken x-axis separates two speed ranges – below (outlined) and above 10 cm/s. Dependence of theta and eta amplitude and frequency on speed above 10 cm/s (**I-L**) and within a 0-10 cm/s range (**M-P**). (**I**) Theta-cycle amplitude is similarly correlated with speed in both RW (0.20 ± 0.005, p<10^−10^) and VR (0.20 ± 0.004, p<10^−10^) across all tetrodes. (**J)** Eta-cycle amplitude is positively correlated with speed VR (0.09 ± 0.004, p<10^−10^), but anti-correlated in RW (−0.11 ± 0.005, p<10^−10^) with significant difference between RW and VR (p<10^−10^, *X*^2^ = 676.44). (**K**) Theta-cycle frequency and speed showed significant correlation in RW (n = 991, 0.16 ± 0.004, p<10^−10^), but not in VR (n = 1581, −0.0079 ± 0.0022, P = 0.002) with significant difference between RW and VR (p<10^−10^, *X*^2^ = 604.22). (**L**) No significant correlation between eta frequency and speed in both RW (5.42e-04 ± 0.0035, p = 0.8) and VR (−0.023 ± 0.0024, p = 0.0022). (**M**) Theta-cycle amplitude is similarly correlated with speed in both RW (0.044±0.0023, p<10^−10^) and VR (0.024 ± 0.002, p<10^−10^) across the tetrodes. (**N**) Eta-cycle amplitude is negatively correlated with speed in VR (−0.09 ± 0.003, p<10^−10^), but not in RW 0.0022 ± 0.003, p = 0.1) with significant difference between RW and VR (p<10^−10^, *X*^2^ = 654.93). (**O**) Significant anti-correlation between theta frequency and speed were seen in VR (n = 1581, −0.079 ± 0.0022, p = 0), but not in RW (n = 991, 0.012 ± 0.0019, p<10^−10^) with significant difference between RW and VR (p<10^−10^, *X*^2^ = 920.35). (**P**) No significant correlation between eta frequency and speed in both RW (−0.01 ± 0.0015, p<10^−10^) and VR (−0.023 ± 0.0024, p<10^−10^) (***, P<0.001).

**Fig. S6.**
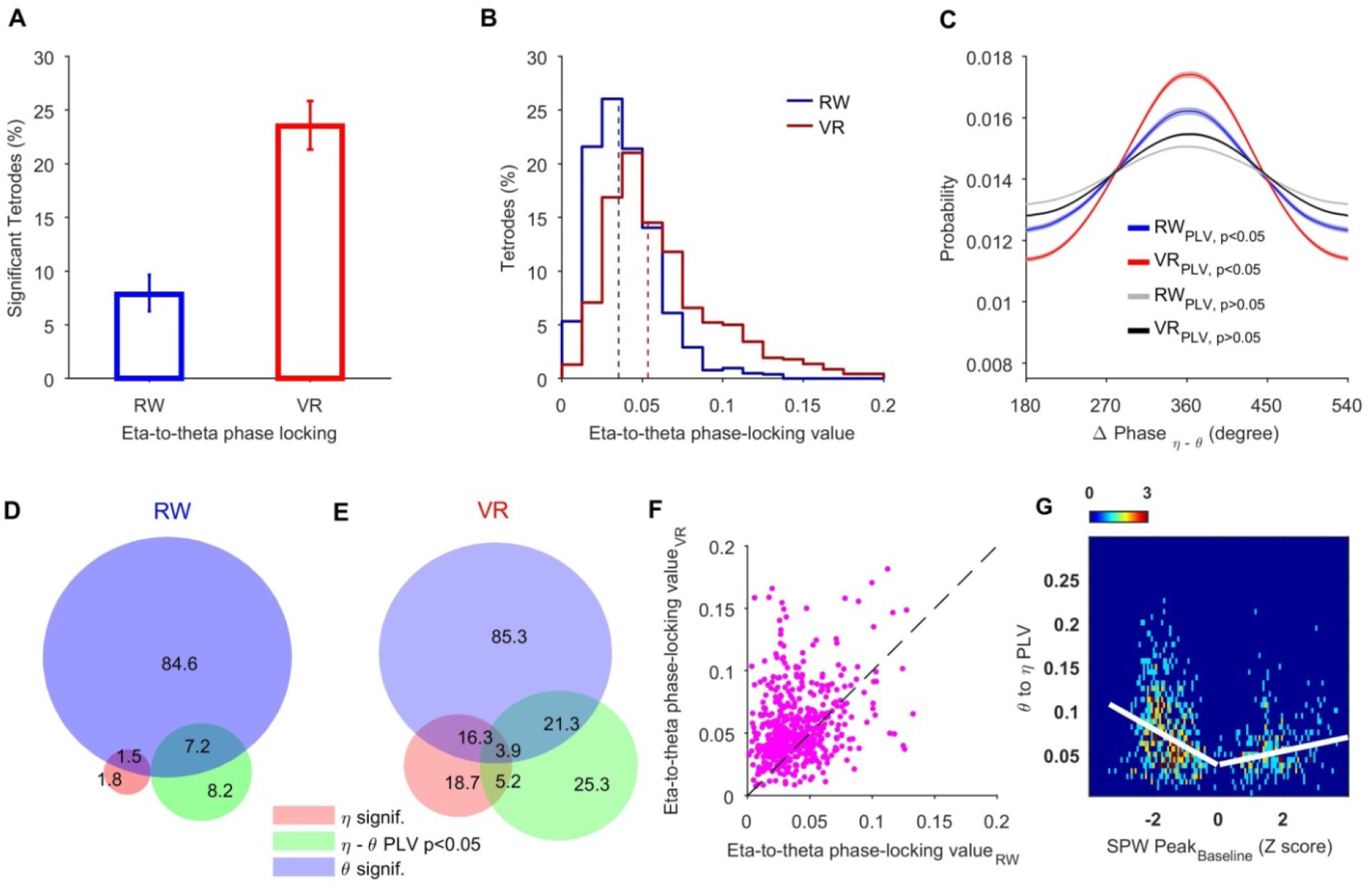
Eta-theta phase-phase coupling is far greater in VR than in RW. **(A)**7.84% (n = 81) and 23.51% (n = 329) of tetrodes showed significant LFP eta-to-theta phase-phase coupling in RW (95% confidence interval [6.28%-9.65%]) and VR (95% confidence interval [21.32%-25.83%]) respectively. The significance of the phase coupling was computed with respect to shuffled data (*n_shuffled_ = 30*) using inverse Fourier transform to randomize phases in time series (α=0.05). (**B)** Phase locking values (PLV) were computed as the mean vector length of the differences between instantaneous LFP theta and eta phases (57, 58). The distribution of the LFP eta-to-theta PLV across the tetrodes was significantly smaller (p<10^−10^, *X*^2^ = 331.56) in RW (0.039 ± 0.0006) than in VR (0.063±0.0017). (**C**) Distributions of eta-to-theta phase differences in RW (blue) and VR (red) for tetrodes with significant PLV. The distributions of non-significant tetrodes are shown in RW (grey) and VR (black). (**D**) Venn diagram of the LFPs with significant theta, eta bands and phase coupling between them in RW. Percentage of LFPs with significant LFP theta (84.6%, n = 838, green), significant eta (1.8%, n = 18, red) and significant eta-to-theta phase locking value (8.2%, n = 81, blue) are given. The intersection of the three circles represents overlapping among the three datasets. (**E**) Similar to **D**, but for VR dataset. Percentage of LFPs with significant theta (85.3%, n = 1042, green), significant eta (18.7%, n = 229, red) and significant eta-to-theta phase locking value (25.3%, n = 309, blue) are given. (**F)** Eta-to-theta PLV for the same tetrodes in RW versus in VR recorded in the same day sessions, showed that 72% of tetrodes had greater eta-theta PLV in VR than RW. (**G**) Relationship between SPW amplitude and polarity and eta-to-theta PLV in VR (for positive SPW n=279, r = 0.18 p = 0.005, for negative SPW, n=617, r = −0.3128, p<10^−10^). Eta-theta phase-phase coupling is larger for tetrodes with larger magnitude SPW, for both +ve and −ve polarity SPW.

**Fig. S7.**
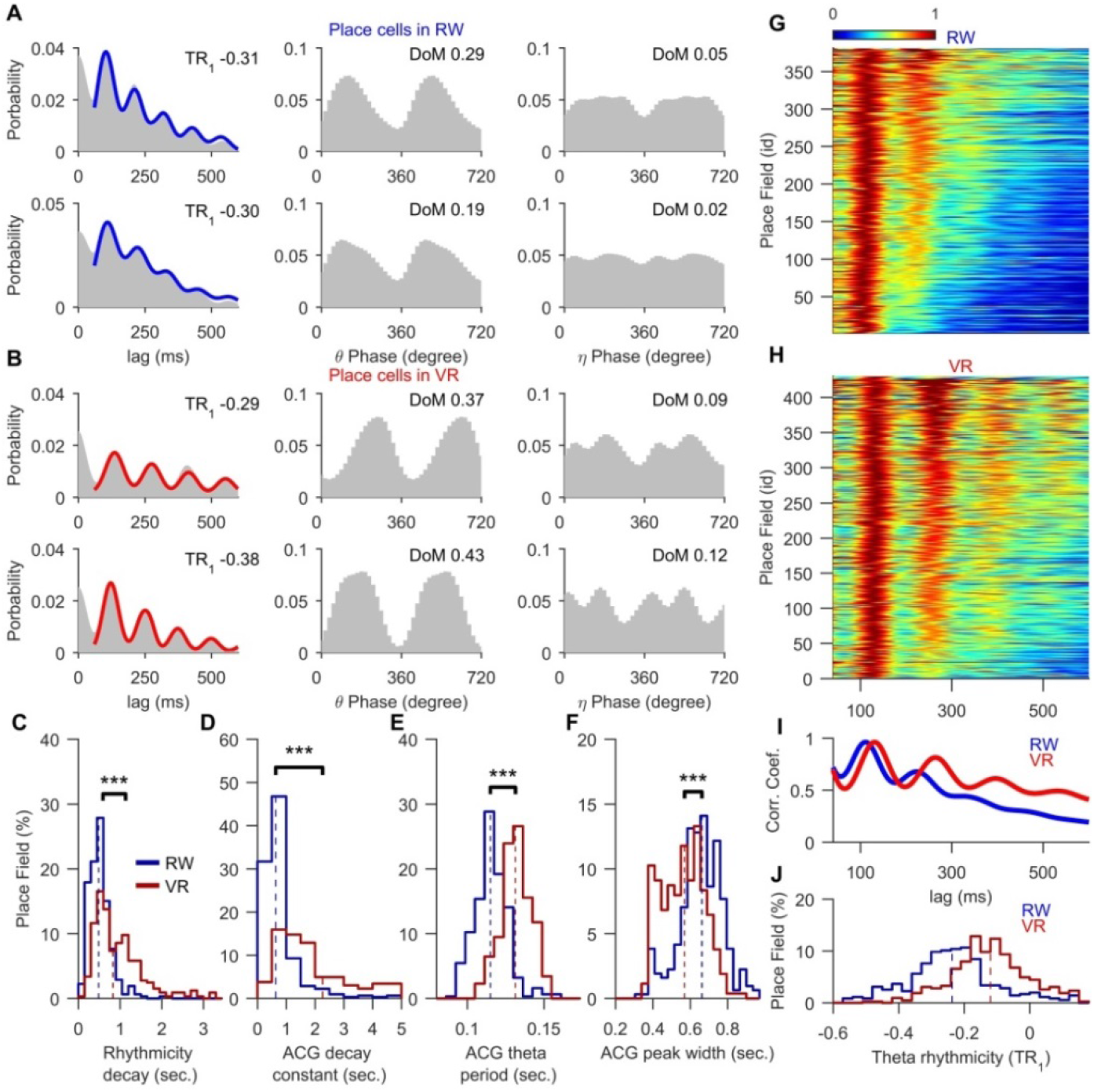
Model based estimate of theta rhythmicity of place fields in RW and VR. **(A, B)** Examples of place cell ACG (grey shaded area) along with fits using a Gaussian mixture model (GMM, see methods) in RW (**A**, left column, blue) and in VR (**B**, left column, red). The theta (middle column) and eta (right column) phase distributions for these example place fields. (**C)** Histograms of ACG rhythmicity decay in RW (blue, 0.59± 0.0439, n = 312) and VR (red, 1.13±0.059, n = 326 are significantly different (p<10^−10^, χ² = 107.92). Thus, ACG in VR decayed nearly half as much as RW. (**D)** Histograms of ACG decay constant in RW (blue, 1.06± 0.08) and VR (red, 3.81± 0.16) show that the ACG decayed more than twice 785 as slowly than in RW (p<10^−10^, χ² = 427.37) indicating sustained theta rhythmicity. (**E**) Histograms of ACG theta period in VR (0.13±0.0006 sec.) is significantly greater (p<10^−10^, χ² = 427.38) than in RW (0.11±0.0006 sec.). (**F**) Histograms of ACG theta peak widths in VR (0.56±0.0065 sec.) are significantly smaller (p<10^−10^, χ² = 89.37) than in RW (0.66±0.007 sec.), showing greater theta rhythmicity. (**G**, **H)** Heat maps of the GMM estimates of ACGs, sorted by increasing TR_1_ for the place fields recorded during running in RW (**G**) and VR (**H**). The ACGs are normalized by their first theta peak values for easy comparison. (**I)** Population average of ACGs show greater theta rhythmicity in VR than in RW. (**J)** Histograms of GMM corrected estimates of the TR1 distributions show VR (median = −0.12) is 75% greater than RW (median= −0.21) (p < 10^−10^, *X*^2^ = 135.54).

**Fig. S8.**
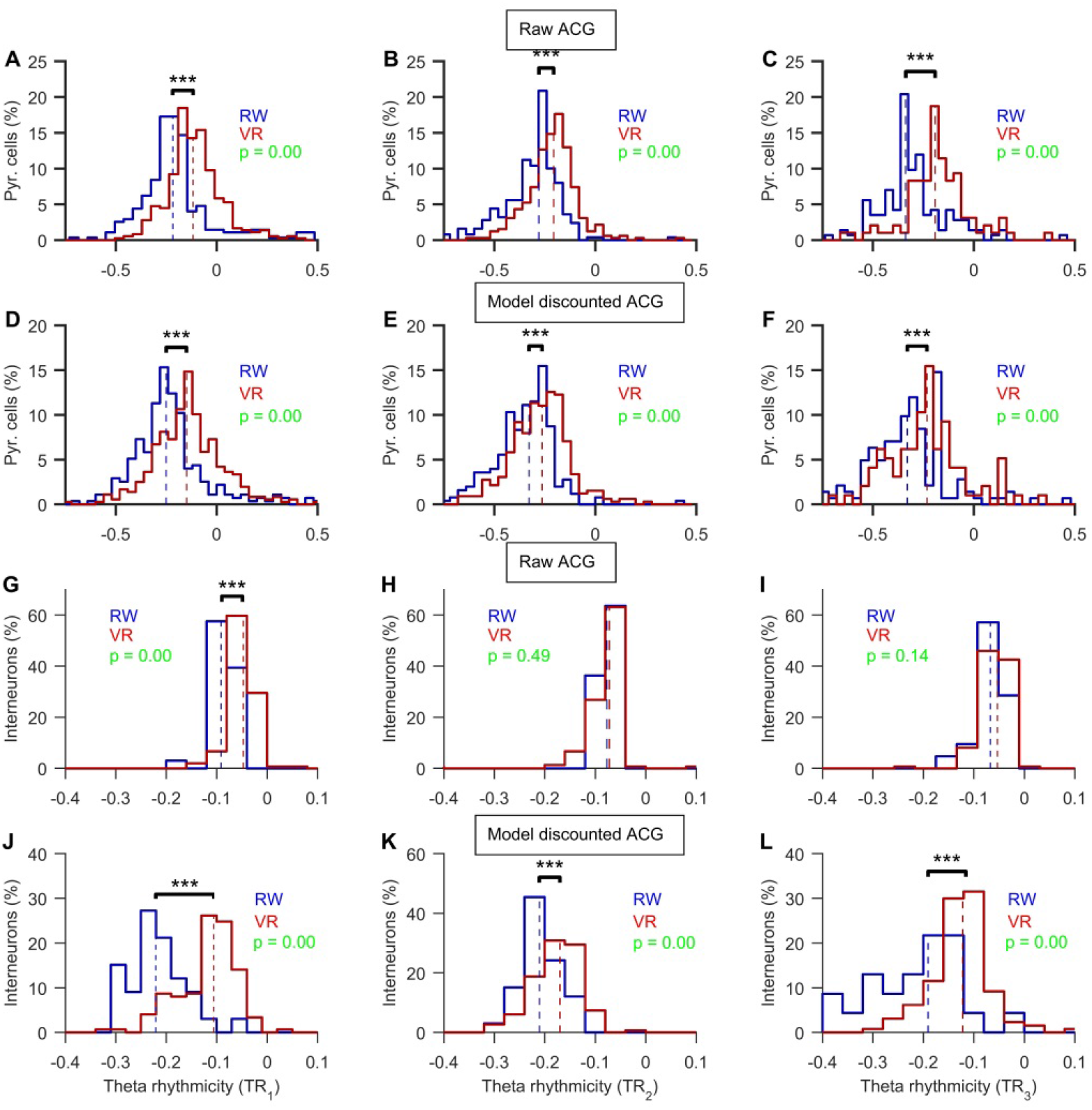
Theta rhythmicity index of the putative pyramidal cells and interneurons in RW and VR. **(A-F)** Data from putative pyramidal cells. **(A)** TR_1_ distributions in VR (−0.1±0.0075, n = 355) is much greater (p < 10^−10^, χ2 = 72.83) than RW (−0.17±0.014, n= 268) and. **(B)** Difference of third-to-second peak of theta (TR_2_) in VR (− 0.21±0.007, n = 153) is much greater (p < 10^−10^, χ2 = 51.83) than RW (−0.32± 0.0081, n= 186). **(C)** Difference of fourth-to-third theta peak (TR_3_) in VR (−0.21±0.008, n = 201) is much greater (p < 10^−9^, χ2 = 33.58) than in RW (−0.36±0.013, n= 114) and. **(D)** Same as a, but model corrected ACG estimates show TR_1_ in VR (− 0.14±0.0091, n = 357) is much greater (p < 10^−10^, χ2 = 54.44) than in RW (−0.21±0.014, n=274) and. **(E)** Difference of third-to-second peak (TR_2_) difference index (p = 2.5188e-05, χ2 = 17.75) between RW (− 0.32±0.0153, n=252) and VR (−0.276±0.009, n =155). **(F)** Difference of fourth-to-third peak (TR_3_) difference index (p = 0.002, χ2 = 9.58) between RW (−0.35±0.014, n=142) and VR (−0.27±0.009, n =76). (**G-L)** Similar to A-D but for interneurons**. (G**) Significant difference of TR_1_ distributions (p < 10^−10^, χ2 = 47.56) cells between RW (−0.09±0.004, n= 33) and VR (−0.05±0.0033, n = 149). (**H**) Difference of third-to-second peak (TR_2_) difference index (p = 0.49, χ2 = 0.47) between RW (−0.0767±0.0031, n= 33) and VR (−0.078±0.0035, n = 149). (**I)** Difference of fourth-to-third peak (TR3) difference index (p = 0.13, χ2 = 2.18) between RW (−0.066±0.006, n= 33) and VR (−0.05±0.007, n = 148). (**J**) Significant difference of TR_1_ distributions (p = 4.8071e-12, χ2 = 47.76) of putative pyramidal cells between RW (−0.215±0.0097, n=33) and VR (−0.116±0.0048, n = 149). (**K)** Difference of third-to-second peak (TR_2_) difference index (p < 10^−5^, χ2 = 16.29) between RW (−0.21±0.0075, n=33) and VR (−0.153±0.0098, n =149). (**L)** Difference of fourth-to-third peak (TR_3_) difference index (p = 2.3933e-06, χ2 = 22.25) between RW (−0.21±0.0187, n=23) and VR (−0.115± 0.0076, n =130). (***, P<0.001).

**Fig. S9.**
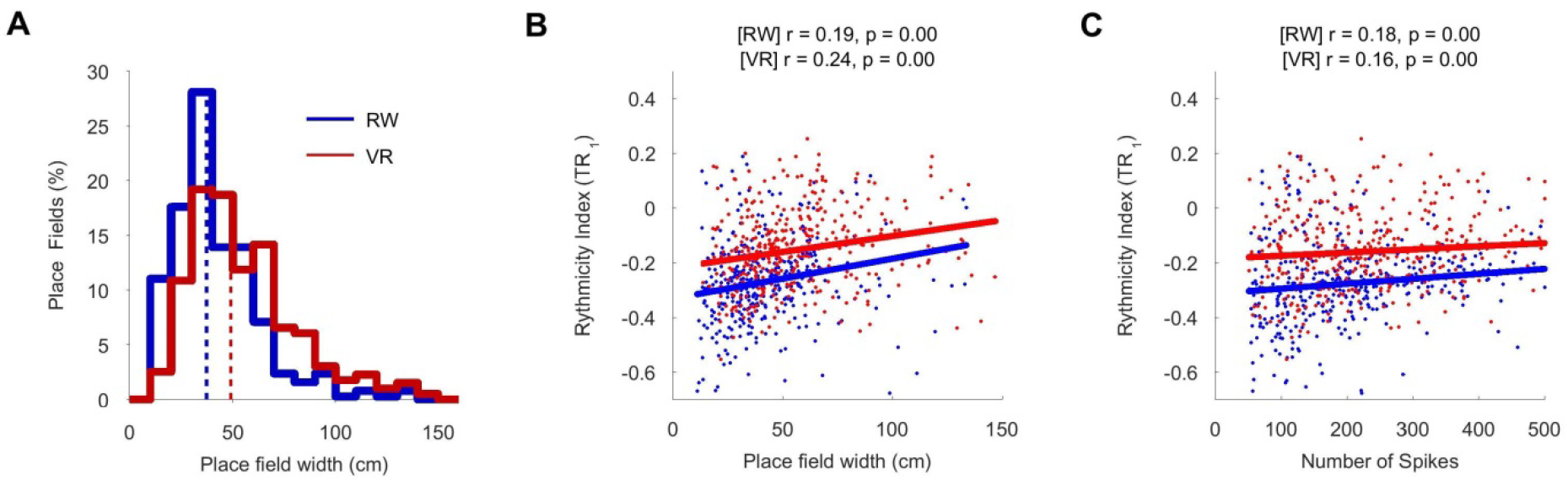
Theta rhythmicity is greater in VR than RW even when factoring out place field width and number of spike contribution. **(A)** Place fields are broader (p < 10^−10^, χ2 = 64.1158) in VR (55.3±1.27cm) than in RW (42.2±1.08 cm). **(B)** Theta rhythmicity index TR_1_ increases as a function of the place field width in RW (blue, r = 0.34, p < 10^−10^) and VR (red, r = 0.33, p < 10^−10^). Linear regression fits are shown. TR1 is consistently greater in VR than in RW across all place field widths. **(C)** TR_1_ increases with the number of spikes in place field, in RW (blue, r = 0.22, p < 10^−10^) and VR (red, r = 0.20, p < 10^−5^). Linear regression fits are shown (blue - RW, red - VR).

**Fig. S10.**
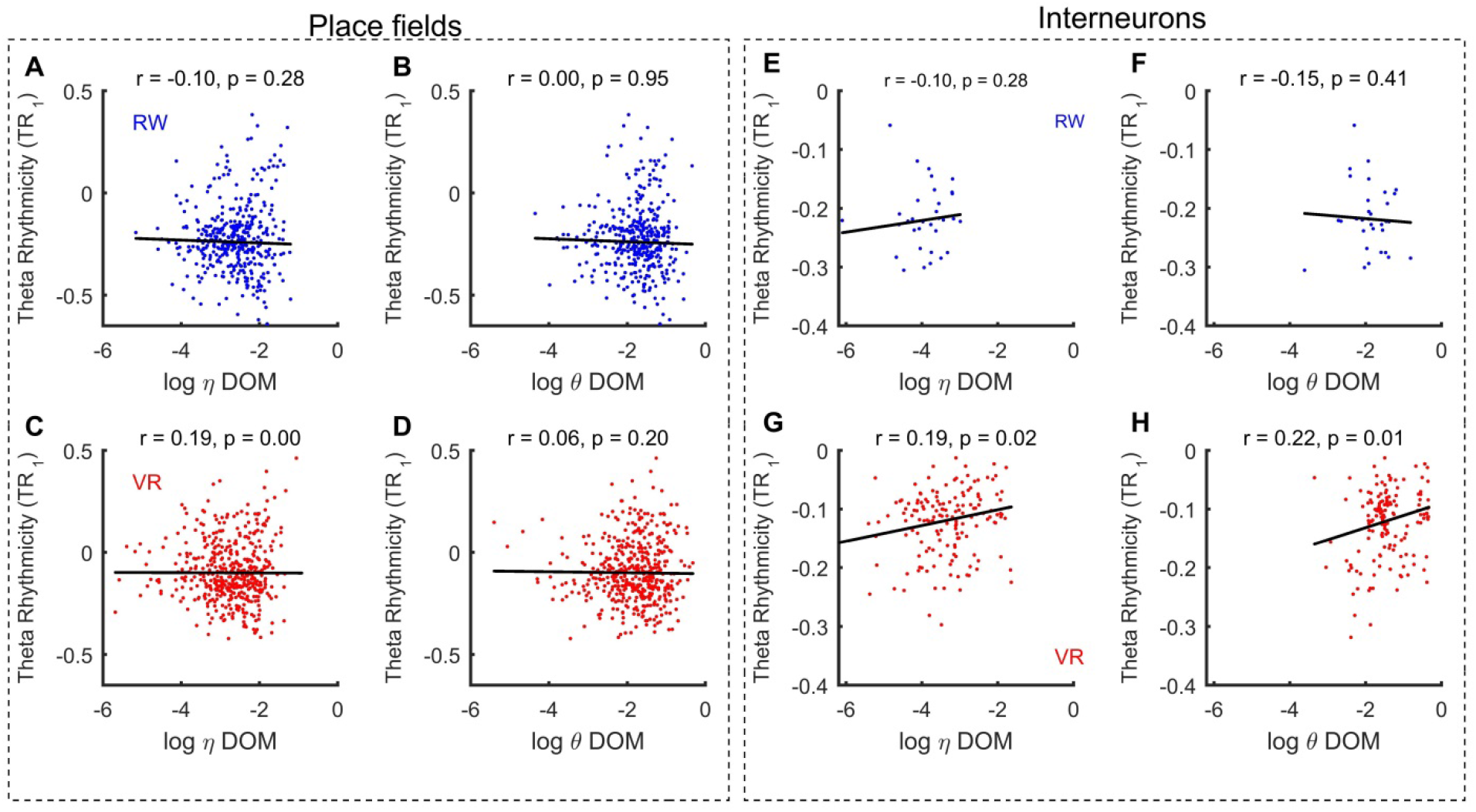
Relationship between theta rhythmicity and theta and eta phase locking of place cells and interneurons in RW and VR. **(A, B)** We quantified the relationship between TR_1_ and eta (r = −0.1, p =0.28, partial Pearson correlation with number of spikes as controlling variable) and theta (r = 0.0032, p =0.95) phase locking of place cells in RW. **(C, D)** The place cells with higher TR_1_ showed increasingly more eta (r = 0.22, p < 10^−5^), but not theta (r = 0.06, p = 0.2), phase locking in VR. **(E, F)** No systematic relationship was found between TR_1_ and (**E**) eta (r = −0.1, p =0.28) or (**F**) theta (r = −0.15, p =0.41) phase locking in RW for interneurons. **(G, H)** Instead, interneurons with higher TR_1_ showed increasingly more (**G**) eta (r = 0.19, p = 0.02) and (**H**) theta (r = 0.22, p =0.01) phase locking in VR.

**Fig. S11.**
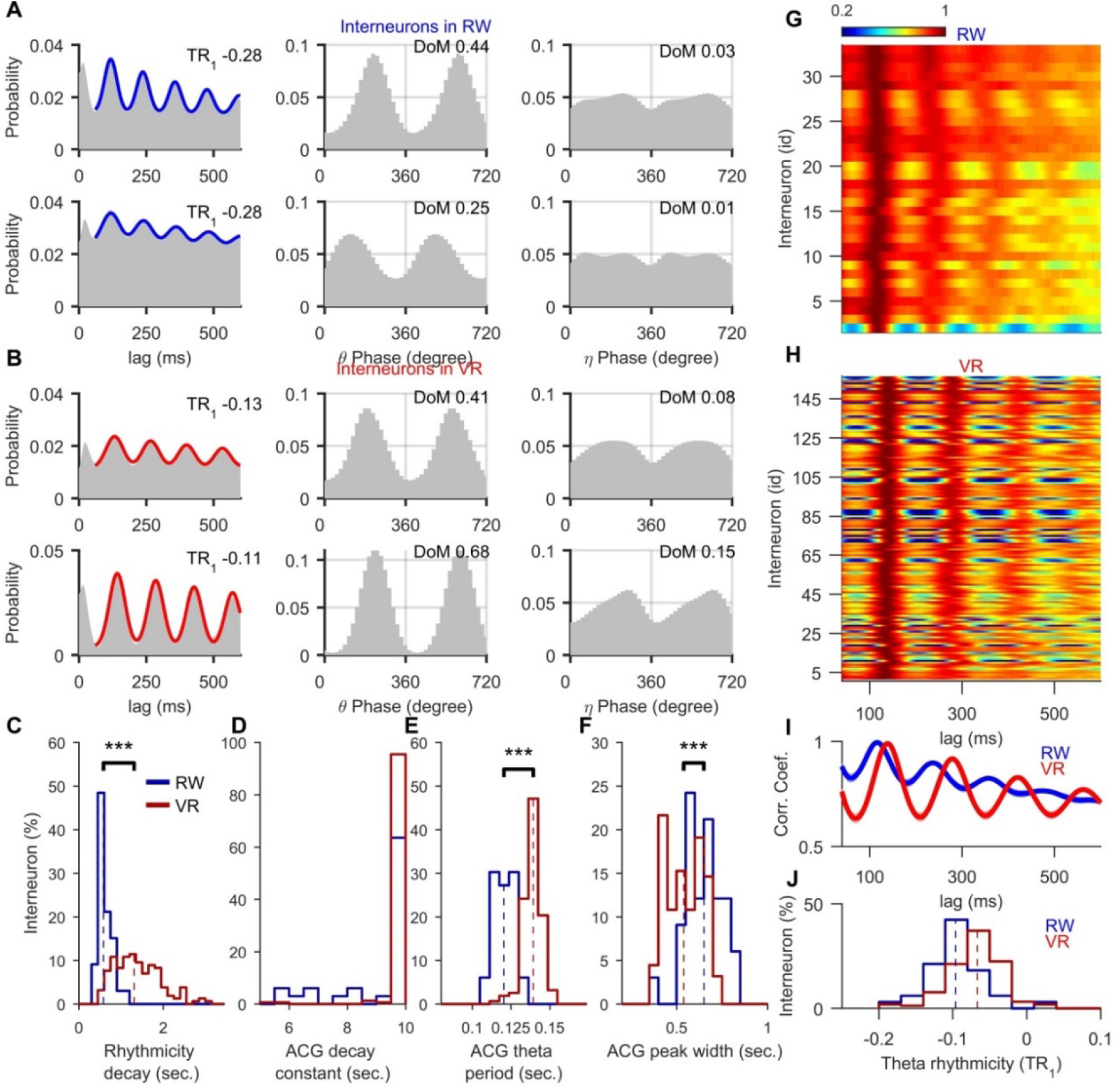
Model fit of autocorrelograms of interneurons in RW and VR. **(A, B)** Examples of interneurons’ autocorrelograms (grey) with TR_1_ values are shown along with fits using GMM in RW (**A**, top two rows, left, blue) and in VR (**B**, bottom two rows, left, red). The distribution of spikes’ theta (middle column) and eta (right column) phases are given. **(C)** Histograms of ACG rhythmicity decay in RW (blue, n = 36, 0.64 ± 0.04) and VR (red, n = 157, 1.4 ± 0.04) are significantly different (p < 10^−10^, χ² = 81.37). **(D)** Histograms of ACG decay constant in RW (blue, 8.4± 0.39) and VR (red, 9.8±0.07) are shown (p < 10^−5^, χ² = 19.65). **(E)** Histograms of ACG theta period in RW (0.12± 0.015) and VR (0.138± 0.02) are significantly different (p < 10^−10^, χ² = 82.63). **(F)** Histograms of ACG peak widths in RW (0.65± 0.002) and VR (0.54± 0.0012) are significantly different (p = 1.1*10^−7^, χ² = 23.71). **(G, H)** Heat map of ACGs of spike trains sorted by increasing TR_1_ for putative interneurons recorded during running in RW (**G**) and VR (**H**). The ACGs are normalized by their first theta peak values. **(I)** The population average of autocorrelations shows greater theta rhythmicity in VR than in RW. **(J)** The histograms of the TR_1_ distributions show significant difference between RW (median = −0.09) and VR (median = −0.07) (p < 10^−10^, χ² = 57.02).

**Table S1.**
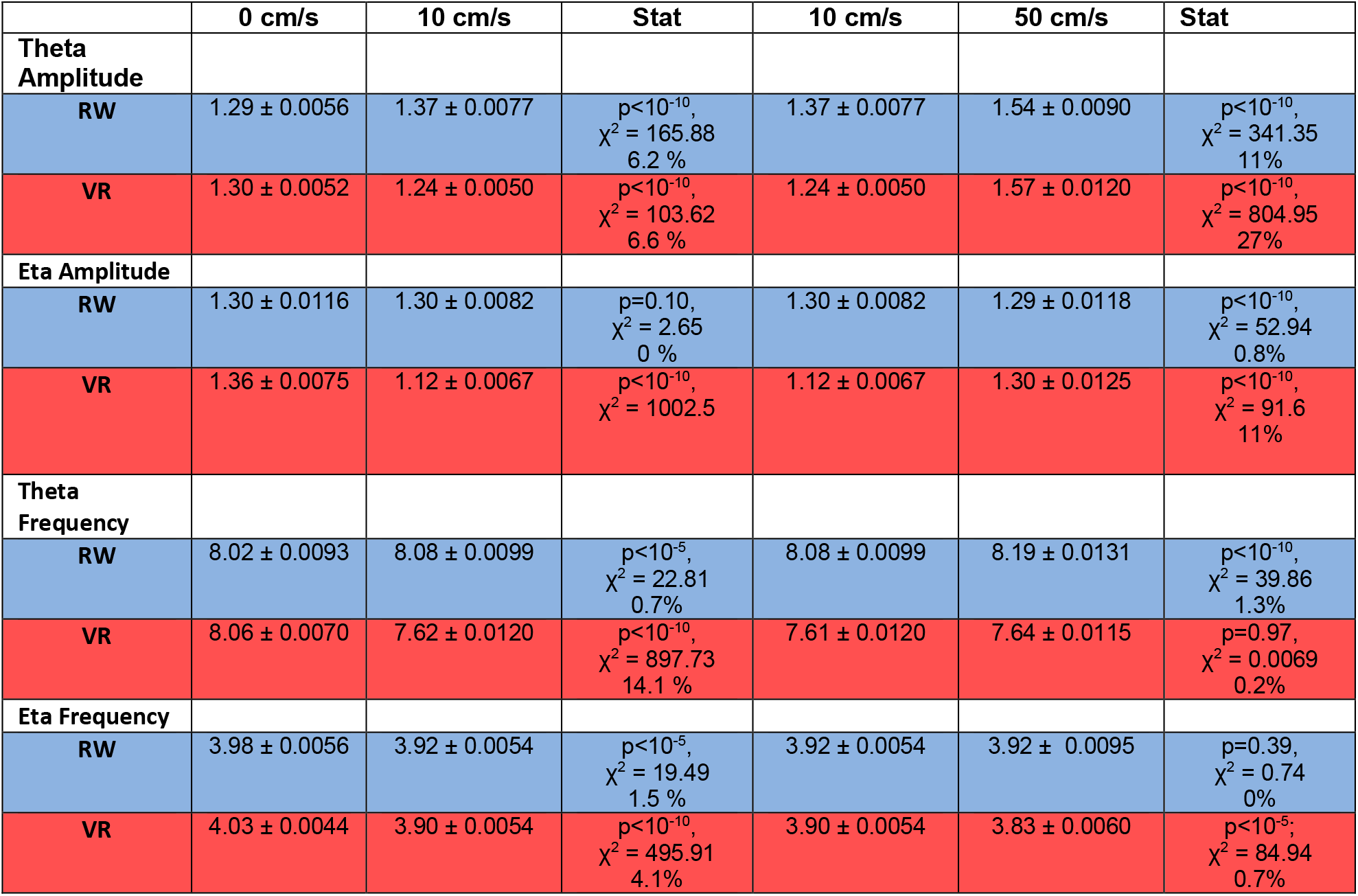
Changes of eta and theta amplitude and frequency in RW and VR at low and high speeds. Population averaged theta and eta amplitudes and frequencies are shown (mean ± s.e.m.) are given in RW (blue, n = 991) and VR (red, n = 1579) at 0, 10 and 50 cm/s. Stat provides significance test using non-parametric Kruskal-Wallis test.

|  | 0 cm/s | 10 cm/s | Stat | 10 cm/s | 50 cm/s | Stat |
| --- | --- | --- | --- | --- | --- | --- |
| **Theta Amplitude** |  |  |  |  |  |  |
| **RW** | 1.29 ± 0.0056 | 1.37 ± 0.0077 | p<10<sup>-10</sup>,<br>χ<sup>2</sup> = 165.88<br>6.2 % | 1.37 ± 0.0077 | 1.54 ± 0.0090 | p<10<sup>-10</sup>,<br>χ<sup>2</sup> = 341.35<br>11% |
| **VR** | 1.30 ± 0.0052 | 1.24 ± 0.0050 | p<10<sup>-10</sup>,<br>χ<sup>2</sup> = 103.62<br>6.6 % | 1.24 ± 0.0050 | 1.57 ± 0.0120 | p<10<sup>-10</sup>,<br>χ<sup>2</sup> = 804.95<br>27% |
| **Eta Amplitude** |  |  |  |  |  |  |
| **RW** | 1.30 ± 0.0116 | 1.30 ± 0.0082 | p=0.10,<br>χ<sup>2</sup> = 2.65<br>0 % | 1.30 ± 0.0082 | 1.29 ± 0.0118 | p<10<sup>-10</sup>,<br>χ<sup>2</sup> = 52.94<br>0.8% |
| **VR** | 1.36 ± 0.0075 | 1.12 ± 0.0067 | p<10<sup>-10</sup>,<br>χ<sup>2</sup> = 1002.5 | 1.12 ± 0.0067 | 1.30 ± 0.0125 | p<10<sup>-10</sup>,<br>χ<sup>2</sup> = 91.6<br>11% |
| **Theta Frequency** |  |  |  |  |  |  |
| **RW** | 8.02 ± 0.0093 | 8.08 ± 0.0099 | p<10<sup>-5</sup>,<br>χ<sup>2</sup> = 22.81<br>0.7% | 8.08 ± 0.0099 | 8.19 ± 0.0131 | p<10<sup>-10</sup>,<br>χ<sup>2</sup> = 39.86<br>1.3% |
| **VR** | 8.06 ± 0.0070 | 7.62 ± 0.0120 | p<10<sup>-10</sup>,<br>χ<sup>2</sup> = 897.73<br>14.1 % | 7.61 ± 0.0120 | 7.64 ± 0.0115 | p=0.97,<br>χ<sup>2</sup> = 0.0069<br>0.2% |
| **Eta Frequency** |  |  |  |  |  |  |
| **RW** | 3.98 ± 0.0056 | 3.92 ± 0.0054 | p<10<sup>-5</sup>,<br>χ<sup>2</sup> = 19.49<br>1.5 % | 3.92 ± 0.0054 | 3.92 ± 0.0095 | p=0.39,<br>χ<sup>2</sup> = 0.74<br>0% |
| **VR** | 4.03 ± 0.0044 | 3.90 ± 0.0054 | p<10<sup>-10</sup>,<br>χ<sup>2</sup> = 495.91<br>4.1% | 3.90 ± 0.0054 | 3.83 ± 0.0060 | p<10<sup>-5</sup>,<br>χ<sup>2</sup> = 84.94<br>0.7% |

